# 3D Epigenome Evolution Underlies Divergent Gene Regulatory Programs in Primate Neural Development

**DOI:** 10.1101/2025.03.11.642620

**Authors:** Silvia Vangelisti, Faye Chong, Tamina Dietl, Vera Zywitza, Sebastian Diecke, Wolfgang Enard, Boyan Bonev

**Affiliations:** Pioneer Campus, Helmholtz Center Munich, 85764 Neuherberg, Germany; Physiological Genomics, Biomedical Center, Ludwig-Maximilians-Universität München, Munich, Germany; Pluripotent Stem Cells Platform, Max Delbrück Center for Molecular Medicine in the Helmholtz Association (MDC), Berlin, Germany; Anthropology and Human Genomics, Faculty of Biology, Ludwig-Maximilians Universität München, 82152 Munich, Germany

**Keywords:** Neocortex expansion, primate neurogenesis, 3D epigenome evolution, gene regulatory programs, comparative genomics

## Abstract

The expansion of the neocortex is a hallmark of human evolution and is closely linked to neural stem cell biology. Yet, the epigenetic mechanisms driving divergent gene regulation during primate neurogenesis remain elusive. Here, we comprehensively mapped 3D genome organization, chromatin accessibility and gene expression in induced pluripotent stem cells and derived neural stem cells from human, chimpanzee, gorilla and macaque. We identified human-specific epigenetic signatures including cis-regulatory regions and enhancer-promoter interactions and linked them to gene regulatory dynamics. Deep learning models revealed that complex regulatory grammar at cis-regulatory regions, including transcription factor binding sites, local context and higher-order chromatin organization, underlies species and cell type-specific differences. High-resolution Hi-C uncovered unexpected global shifts in 3D genome architecture in chimpanzee and gorilla neural stem cells while topologically associating domains remain remarkably conserved. Notably, species-specific genes interacted with multiple differentially accessible regions, suggesting that synergistic enhancer activation is a key mechanism driving epigenome evolution. These findings provide new insights into the epigenetic basis of primate brain evolution.

## Introduction

The increased size of the neocortex in humans is closely linked to the biology and proliferative capacity of the neural stem/progenitor cells (NSC)^1–4^. While some NSC populations appear to be conserved, notable differences such as cell cycle duration have been identified, particularly in the context of primate evolution^5,6^. Furthermore, in addition to biochemical and metabolic differences, epigenetic regulation has emerged as an important layer for determining fate choices in the developing neocortex and has been closely linked to human brain evolution.

In particular, recent advances in comparative genomic and cellular analyses point to the non-coding genome as a key driver in shaping the evolution of complex human-specific phenotypes^7–15^. For example, cis-regulatory regions (CREs) are associated with rapid turnover rates during evolution^16,17^, and species-specific enhancers are frequently active in narrow spatial and temporal windows compared to conserved elements^9^. Human-specific regulatory changes, such as deletion of the *GADD45G* enhancer^18^, sequence changes in the *FZD8* enhancer^19^, and loss of a transcription factor (TF) motif within the *CBLN2* enhancer^20^, have been shown to affect gene expression and contribute to human neocortex expansion. Furthermore, evolutionary-related changes in the 3D genome organization, such as the emergence of novel topologically associating domains (TADs) and loops^9,21^, rewiring of human accelerated regions (HARs) and their target genes^22^ and enhancer hijacking^23^ can all change the regulatory landscape of human cortical development. Finally, deep learning models have emerged as an important tool for linking changes in the linear genome to cell type- or species-specific enhancer dynamics^9,23–26^, allowing us to discriminate between variations in the DNA sequence and the contribution of the epigenetic context to human brain evolution.

A major limitation is that most studies have focused on a single cell type^10^ or used bulk profiling in organoids/tissues^21^, making it difficult to distinguish species-specific from cell type-specific changes. In addition, many studies have used only two species or compared distantly related species, which complicates the identification of truly human-specific regulatory changes^9,21^. Furthermore, most studies examining 3D genome organization in the context of primate evolution have focused specifically on selected elements such as HARs, making it more difficult to identify global genome-wide changes and relate them to chromatin accessibility and transcription. Broadening the analysis to include closely related species^27,28^, focusing on specific cell types and integrating multiple epigenetic layers promises to fill this gap and provide deeper insights into the molecular basis of the emergence of complex biological processes during evolution.

To gain deeper insights into the epigenetic mechanisms regulating early cortical development during primate evolution, we systematically mapped the transcriptome, chromatin accessibility and 3D genome organization in induced pluripotent stem cells (IPSC) and NSC in four closely related primate species (crab-eating macaque, gorilla, chimpanzee and human). Using these unique datasets, we identified and quantified changes in cell type-specific epigenetic signatures that have emerged during primate evolution, with a focus on those specific to humans. While human-specific structural variants (fhSVs) and HARs are associated with gene expression and epigenomic changes during evolution, they alone do not fully explain these differences. Instead, changes in TF motifs emerge as key elements of species-specific regulatory evolution. We further validated our human-specific enhancer predictions using cortical organoids and adapted deep learning models to dissect epigenome evolution, pointing to complex regulatory grammar at CREs, the local context and potentially the proximity to other genomic elements as contributing to species- and cell type-specific differences. In addition, we confirm the conservation of TADs throughout primate evolution and uncover human-specific regulatory contacts that may have contributed to neocortical expansion. However, these contacts alone are not sufficient to explain the evolution of the primate transcriptome, supporting the notion that the regulation of cellular identity is a complex event involving multiple molecular layers of epigenetic regulation. Taken together, these findings provide new insights into the molecular basis of primate brain evolution.

## Results

To investigate how gene regulatory programs have evolved during primate neocortical development, we employed a comparative epigenomics approach in four closely related primate species: crab-eating macaque (M), gorilla (G), chimpanzee (C) and human (H), and two cellular contexts: induced pluripotent stem cells (IPSC) and neural stem/progenitor cells (NSC) (Figure 1A-B). Using bulk RNA-seq, ATAC-seq, and high-resolution Hi-C, we profiled gene expression, chromatin accessibility, and three-dimensional genome organization during primate neurogenesis, generating one of the most comprehensive datasets of primate epigenome evolution to date (Figure 1C).

**Figure 1.**
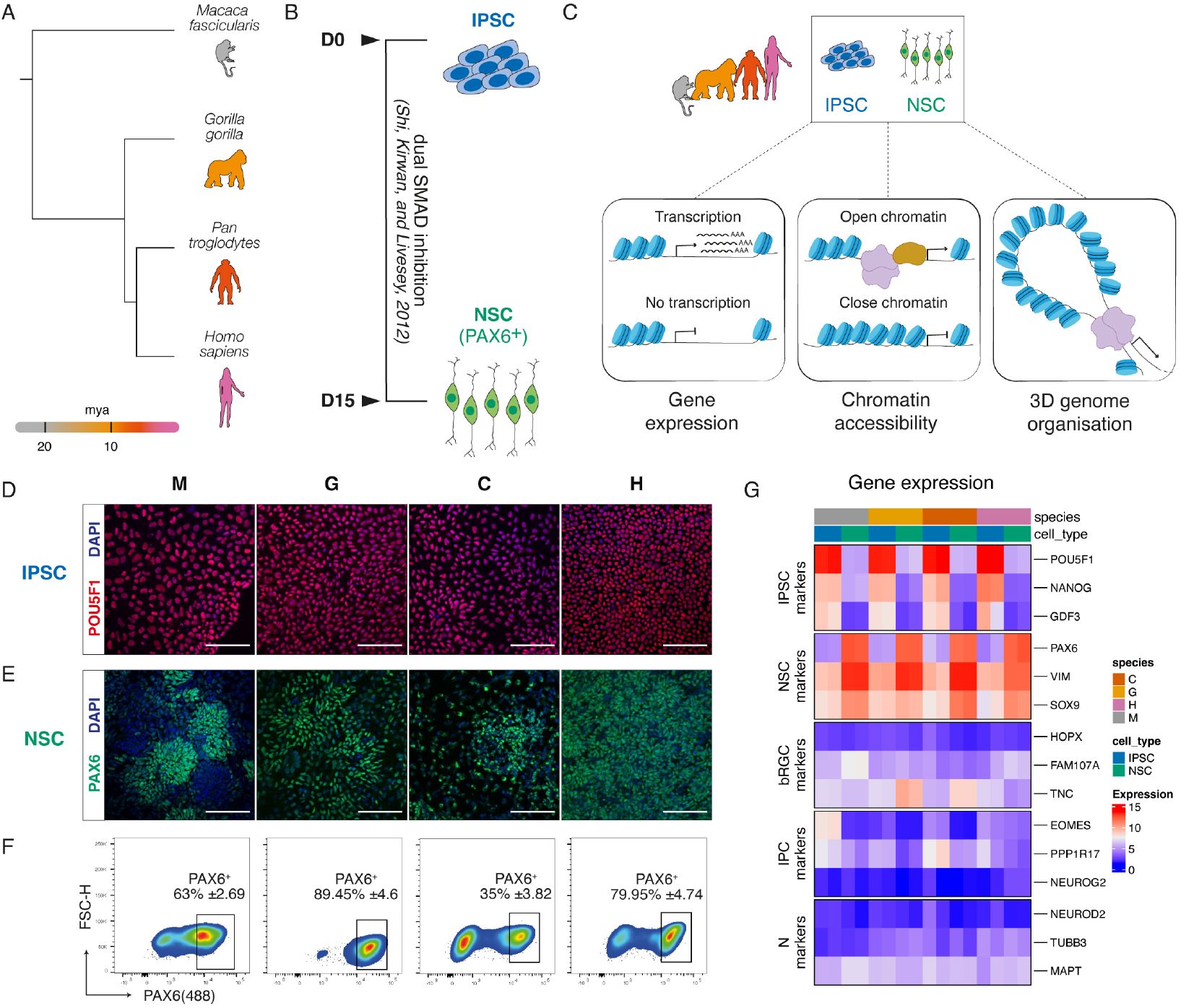
Primate IPSC-derived neural stem cells as a model system to study human brain evolution. (A) Dendrogram representing the evolutionary distance of each species in our study based on^7^. Mya, million years ago. (B) Schematic of dorsal neural progenitor differentiation, showing NSC generation from IPSC. D, day. (C) Overview of the experimental design. (D-E) Representative immunofluorescence images of primate IPSC (D) and NSC at day 15 (E). Scale bar, 100 µm. (F) Gating strategy for isolating PAX6^+^ NSC based on FACS. The numbers represent the mean ± SD from the parental singlets population. (G) Heatmap showing the expression of cell type-specific genes in primate IPSC and NSC. M, crab-eating macaque; G, gorilla; C, chimpanzee; H, human.

We first assessed the quality of the IPSC via karyotyping and immunofluorescence. Overall, they exhibited a normal karyotype, with only very minor aberrations observed (Figure S1A). We further confirmed IPSC quality by staining colonies with established pluripotency markers, including SOX2, NANOG, and POU5F1, prior to each differentiation round (Figure 1D, S1B).

To obtain insights into the convergent and divergent types of gene regulation during neurogenesis, we differentiated the IPSC into cortical NSC using a well-established dual-SMAD inhibition *in vitro* protocol^29,30^ (Figure 1B, 1D–E, S1B–C). To validate our 2D differentiation system, we performed immunostaining at 15 (D15) and 27 days (D27) post-neural induction for the NSC marker PAX6, the intermediate progenitor cell (IPC) marker EOMES, and the pan-neuronal marker TUBB3 (Figure S1C–D). Although NSC across all species were efficiently generated, the proportion of PAX6^+^ cells varied among species (Figure 1E, S1C). To account for this variability, we purified PAX6^+^ NSC via fluorescence-activated cell sorting (FACS), resulting in a homogenous population of NSC for subsequent analysis (Figure 1F).

We further validated the identity of our cellular populations using transcriptomic analysis. Pluripotency markers (e.g., *POU5F1, NANOG*, and *GDF3*) were robustly expressed in IPSC across species, whereas NSC-specific genes (e.g., *PAX6, VIM*, and *SOX9*) were upregulated in all primate NSC. In addition, markers for basal radial glial cells (bRGCs) – including *HOPX, FAM107A*, and *TNC* – as well as IPCs (*EOMES, PPP1R17, NEUROG2*) and neurons (*NEUROD2, TUBB3*, and *MAPT*) were either low or undetectable (Figure 1G).

Collectively, these results validate our experimental system, providing a robust platform for investigating epigenome evolution at multiple molecular levels. Importantly, the inclusion of three great ape species – gorilla, chimpanzee, and human – alongside an outgroup species (crab-eating macaque) enables us to distinguish epigenomic features uniquely specialized in humans from those conserved among great apes.

### Comparative transcriptomics reveals cell type- and species-specific gene expression

To identify conserved and divergent gene regulatory programs, we first focused on differentially expressed genes (DEGs). To overcome limitations due to incomplete gene annotations in non-model organisms, we uniformly mapped the RNA-seq data to the hg38 genome, as recommended for closely related species^31^. This strategy resulted in a clear separation between cell types and improved correlation among replicates compared to conventional analyses based on individual genomes (Figure 2A, S2A-B).

**Figure 2.**
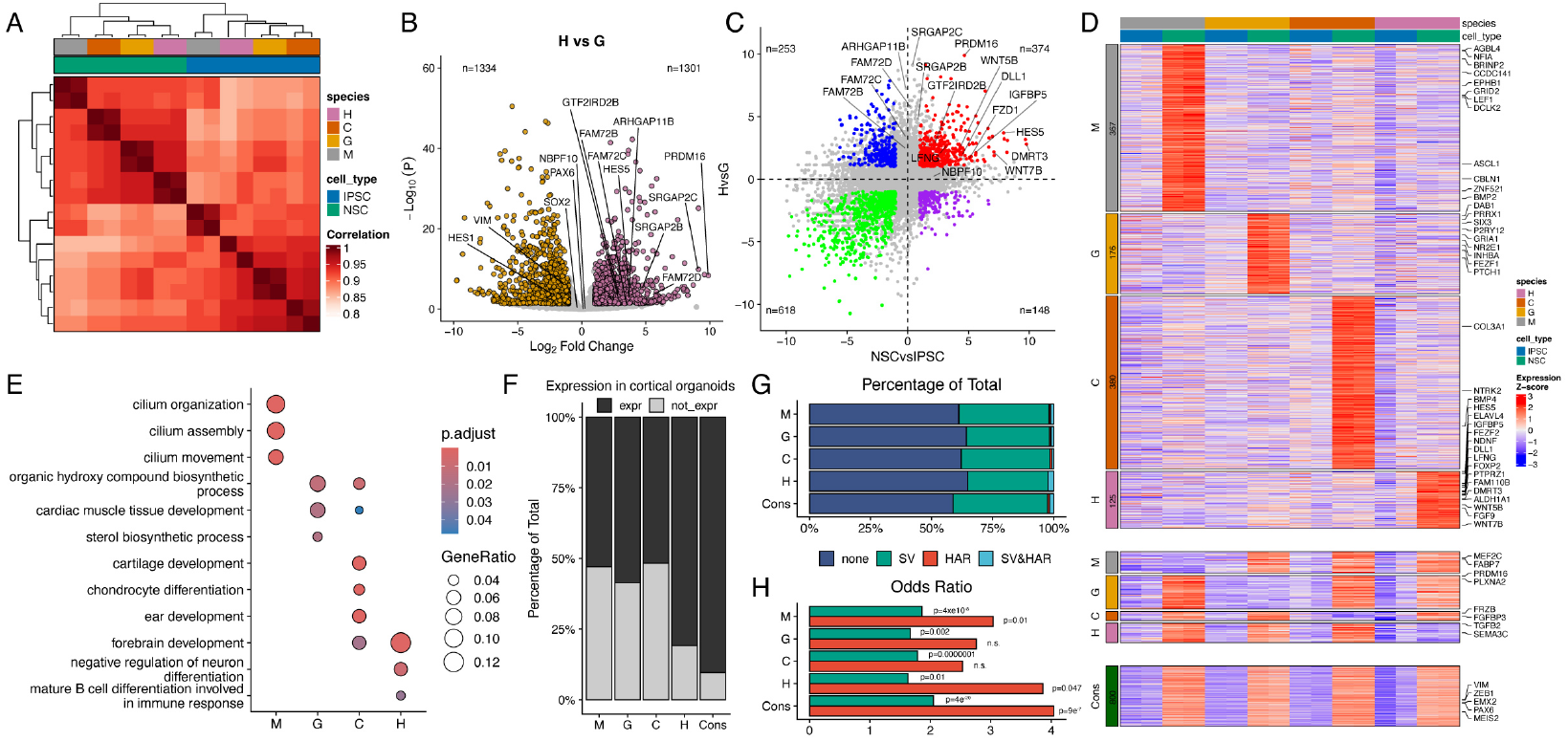
Comparative transcriptomics reveals cell type- and species-specific gene expression. (A) Pairwise correlation matrix displaying Spearman’s correlation coefficient for gene expression (reads mapped to the human genome). (B) Scatterplot depicting differentially expressed genes (DEGs; false discovery rate (FDR) < 0.05) in human vs gorilla NSC. (C) Scatterplot showing a direct four-way comparison of log2 fold changes gene expression in human vs gorilla and NSC vs IPSC. (D) Heatmap depicting significantly upregulated (top), downregulated (middle) and conserved (bottom) primate NSC genes. (E) Dot plot showing the gene ratios of the enriched gene ontology across the upregulated genes in D). The color of the circles represents the Benjamini-Hochberg adjusted P value. (F) Percentage of genes that are expressed (black) or not expressed (grey) in cortical organoids from^44^. (G-H) Percentage and odds ratio of upregulated or conserved genes in D) that overlap with fhSVs^45^ and/or zooHARs^23^. M, crab-eating macaque; G, gorilla; C, chimpanzee; H, human; Cons, conserved.

We next examined gene expression changes between human and gorilla in each cell type independently (Figure 2B, S2C). Consistent with previous studies^32–34^, we confirmed the human-specific expression of several genes (e.g., *GTF2IRD2B, FAM72B, FAM72C, FAM72D, SRGAP2B, SRGAP2C*, and *ARHGAP11B*). However, these genes were not restricted to NSC but were also differentially expressed in human IPSC.

We then asked if we could identify genes that change their expression in both a cell type- and species-specific manner. Surprisingly, most previously identified human-specific genes exhibited human-specific but not cell type-specific expression (Figure 2C). In contrast, we identified 374 novel genes with human-NSC expression, such as the transcription factor PRDM16^35^. Many of them belong to the WNT and NOTCH pathways (e.g., *WNT5B, WNT7B, IGFBP5, DLL1, HES5* and *LFNG*), which are known to be associated with NSC proliferation and differentiation^34,36–39^ and have been implicated in human brain evolution^19,34,39^.

Extending our analysis to include all species and cell types revealed both conserved and species-specific DEGs (Figure 2D, S2D). Among the 125 genes uniquely upregulated in human NSC, we again found key components of the WNT and NOTCH pathways along with novel candidates, such as *DMRT3*, which has been linked to mouse neocortical development^40^, *ALDH1A1* — a key enzyme in retinoic acid biosynthesis implicated in neocortical expansion^20,41^ — and *PTPRZ1*, a gene highly expressed in bRGCs^42^ but not previously associated with human evolution (Figure 2D). Conversely, genes upregulated in human IPSC were associated with FGF signalling, but also included TFs associated with the maintenance of ES cell pluripotency, such as *ZSCAN10* and *ZFP42* (Figure S2D), as well as ion metabolism (Figure S2E).

Genes upregulated in human NSC were associated with Gene Ontology (GO) terms related to forebrain development and negative regulation of neuronal differentiation (Figure 2E), consistent with the extended proliferation phase of NSC in human. Interestingly, macaque-specific NSC genes were associated with cilia organization, which has been linked to changes in cell cycle and neurogenesis in humans^43^.

To validate our findings in another model system, we compared the expression of the different groups of genes in either RGCs or IPCs isolated from human organoids^44^. We observed that the majority of the human NSC-specific genes were also expressed in this system, while NSC genes associated with other species were more frequently repressed (Figure 2F).

Finally, we explored the mechanisms underlying species-specific transcription by examining fixed human-specific structural variants (fhSVs)^45^ and human accelerated regions (HARs)^23^. Although only a small fraction of DEGs were associated with either fhSVs or HARs (Figure 2G, S2F), the overlap between HARs and human-specific genes was significantly enriched in NSC (Figure 2H, S2G). These findings indicate that while HARs contribute to the emergence of human-specific transcriptional profiles, they do not fully account for these differences, as many conserved genes also overlapped with HARs.

Overall, these results identify novel cell type-specific DEGs during primate evolution, emphasizing the importance of considering both cell type and species when interested in genes involved in specific developmental contexts. They also suggest that although SVs and particularly HARs could contribute to evolutionary divergence, they alone may not be sufficient to explain human-specific NSC gene expression.

### Changes in TF motifs during evolution underlie dynamic chromatin accessibility

To investigate the cell type-specific dynamics of the epigenetic landscape during primate evolution, we performed bulk ATAC-seq on each isolated cell population in parallel (Figure 1C). We optimized the protocol^46^ for fixed-sorted cells and confirmed the high quality of the ATAC data (Figure S3A-B).

To identify differentially accessible regions (DARs), we first generated a reference-free Cactus multiple sequence alignment^47^. We then determined contiguous orthologous regions using HALPER^48^ and counted the number of fragments in each replicate, generating a species x regions count matrix. Spearman pairwise correlation revealed a separation mainly driven by cell type differences and a high ATAC correlation between replicates of the same sample (Figure 3A), demonstrating the potential of our approach to identify cell type-specific changes in regulatory regions across species.

**Figure 3.**
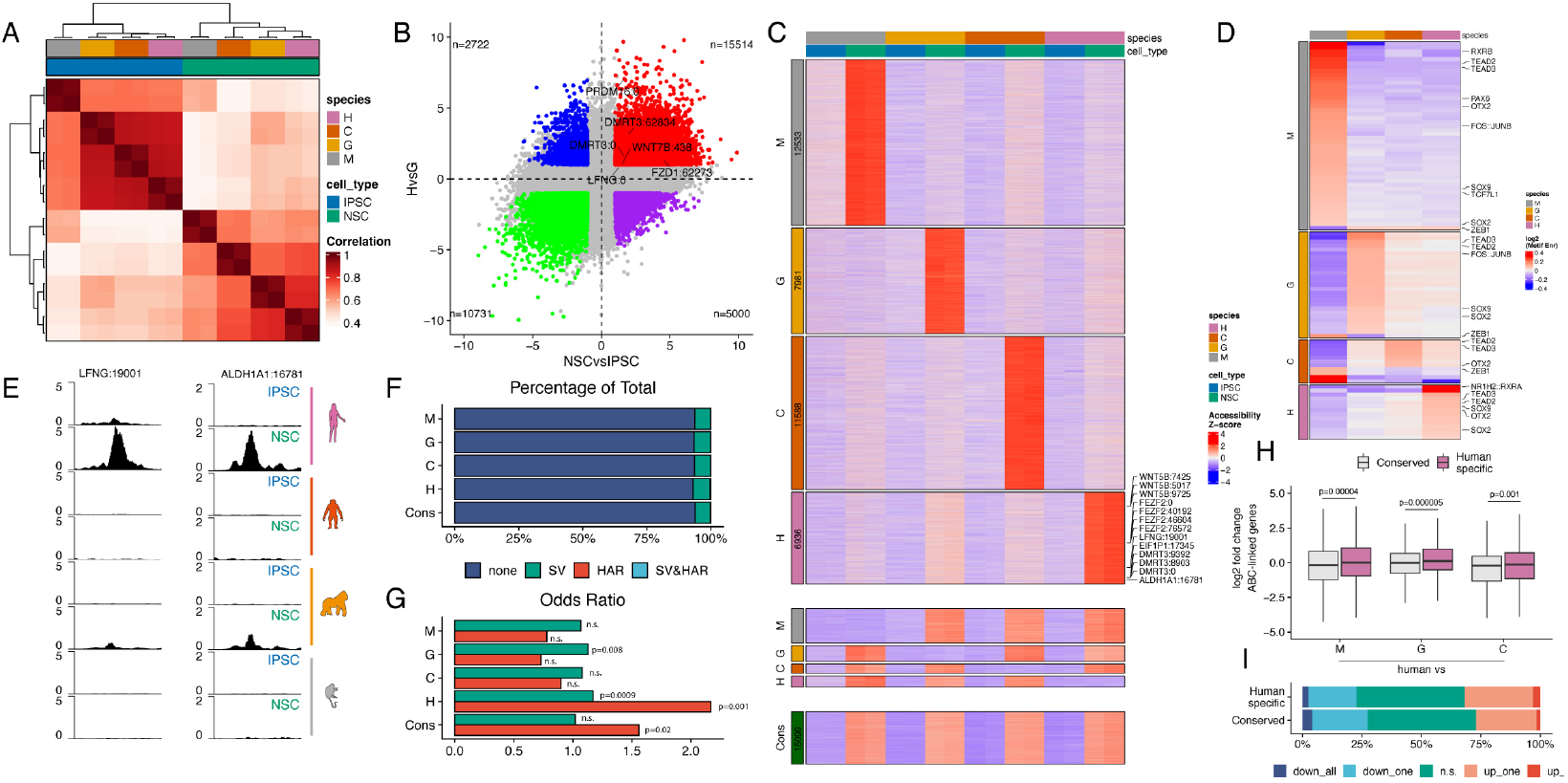
Changes in TF motifs during evolution underlie dynamic chromatin accessibility. (A) Pairwise correlation matrix displaying Spearman’s correlation coefficient for chromatin accessibility. (B) Scatterplot showing a direct four-way comparison of log2 fold changes accessibility in human vs gorilla and NSC vs IPSC. (C) Heatmap depicting differentially accessible regions (DARs) in primate NSC. (D) Heatmap showing TF motif enrichment in the NSC-species specific DARs regions in each species. (E) ATAC-seq tracks of primate IPSC and NSC centred on two human-specific accessible *loci*: *LFNG* (left) and *ALDH1A1* (right). (F-G) Percentage and odds ratio of DARs or conserved regions in C) that overlap with fhSVs^45^ and/or zooHARs^23^. (H) Boxplots displaying gene expression log2 fold change in human compared to NHP for the predicted human NSC-specific DAR target genes (based on the ABC model in human NSC). Statistical significance was calculated using an unpaired two-sided Wilcoxon rank-sum test. (I) Stacked barplot showing the percentage of genes in H) that are downregulated in all species (down_all), downregulated in only one species (down_one), not significant (n.s.), upregulated in one (up_one) or in all species (up_all) when comparing expression in human versus NHP NSC.

First, we compared human and gorilla IPSC and NSC (Figure 3B). We identified 15514 regions that were differentially accessible in human NSC, with some of them associated with the DEGs identified in the previous analysis. We then extended this pairwise comparison to all species and cell types. We identified 6936 human NSC- and 3068 human IPSC-specific regions (Figure 3C, S3C) and confirmed that they are differentially accessible across species and cell types (Figure S3D–E). In addition, epi-conserved elements were also more conserved at the sequence level compared to species-specific NSC DARs (Figure S3I). The human NSC regions were once again associated with proximal or distal CREs close to genes belonging to the WNT and NOTCH pathways (e.g., *WNT5B, FEZF2* and *LFNG*), as well as *DMRT3* and *ALDH1A1* (Figure 3C, 3E).

To identify potential mechanisms underlying the changes in accessibility across species, we performed a TF motif enrichment analysis. We observed that increased accessibility was mainly associated with the gain of TF motifs rather than the loss of existing ones (Figure 3C–D, S3C, S3F). In particular, we found members of the SOX family, FOS::JUNB complex^44,49–51^ and TEAD2/3^51,52^ enriched in all NSC species-specific DARs. In contrast, the motif of the repressor ZEB1 was lost in regions with species-specific accessibility in macaque, gorilla and chimpanzee (Figure 3D). In addition, we found the nuclear receptors NR1H2::RXRA motif enriched in the human open regions, consistent with previous studies highlighting the role of retinoic acid metabolism in human neocortex specification^20,41^. Conversely, we observed that pluripotency-associated TFs such as OCT4 (POU5F1) and SOX2 were enriched in species-specific IPSC regions (Figure S3F). Surprisingly, we also observed that CTCF motifs were also enriched in those sites.

Next, we examined the enrichment of fhSVs and HARs. Our analysis revealed that only a small proportion of DARs overlap with either fhSVs or HARs, regardless of their cell type or species specificity (Figure 3F, S3G). Although small, the overlap between HARs and human-specific DARs was higher than expected by chance, even when we compared it with conserved DARs, specifically in NSC (Figure 3G, S3H). These results suggest that while HARs contribute to human-specific accessibility differences, most of the interspecies accessibility variation is likely driven by other mechanisms.

Transposable elements (TEs) have been previously linked to epigenome evolution^9,53,54^. Therefore, we searched for TEs enriched in the human DARs compared to the epi-conserved regions. We identified a hominid-specific family of retrotransposon elements – SVA^55^ – that was specifically enriched in human NSC DARs but not in IPSC DARs (Figure S3J–K) and has not previously been linked to human brain evolution at the epigenome level.

Finally, to link changes in chromatin accessibility to gene expression, we used an Activity-by-Contact (ABC) model^56^ based on human NSC Hi-C. We found that the predicted target genes of human NSC DARs were upregulated compared to genes linked to conserved DARs (Figure 3H–I). This suggests that although stronger accessibility is overall correlated with increased gene expression, not all DARs are associated with species- and cell type-specific genes.

In summary, these results highlight the power of our approach to identify DARs associated with primate evolution. They also suggest that the emergence of novel TF motifs and SVA TEs is one of the main mechanisms underlying cell type-specific changes in accessibility across species, linking sequence to epigenome evolution. Finally, dynamic CREs are correlated with changes in gene expression but are not sufficient to fully predict the magnitude of the effect.

### Validation of the human-specific NSC DARs in human cortical organoids

To further validate the activity of the human NSC DARs, we decided to use cerebral organoids. We purified radial glia cells (RGCs) and intermediate progenitor cells (IPCs) from day 45 (D45) cortical organoids as previously described^44^ and performed ATAC-seq and Hi-C (Figure 4A, S4A). We found that 42.8% (2971/6936) of the human NSC DARs overlapped with a peak in either RGCs or IPCs, which was significantly higher than other species-specific NSC peaks, such as chimpanzee (9.5%) and was even higher than conserved NSC peaks (35.7%).

**Figure 4.**
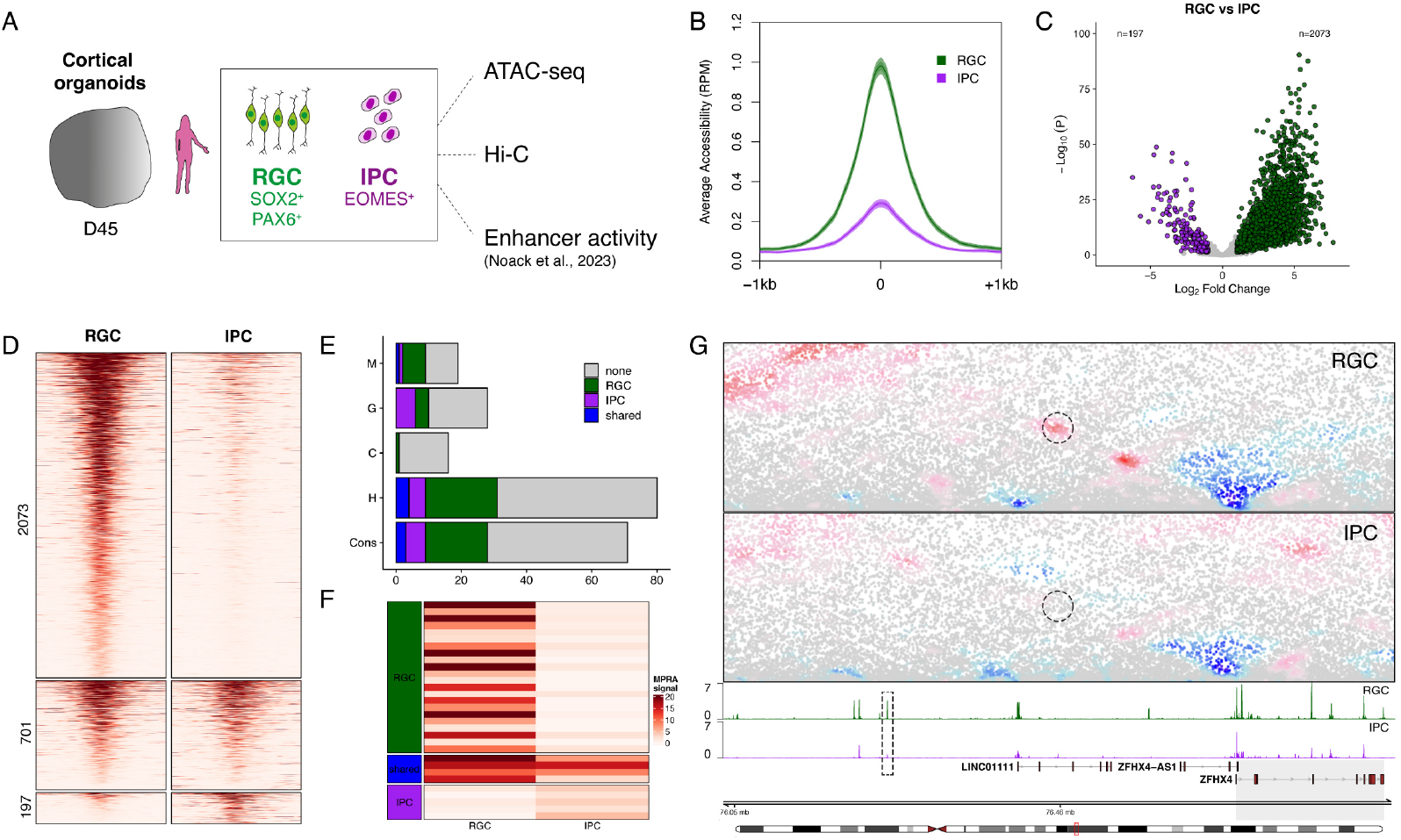
Validation of the human-specific NSC DARs in human cortical organoids. (A) Schematic representation of the experimental approach to isolate radial glia cells (RGCs) and intermediate progenitor cells (IPCs) from cortical organoids. (B) Average accessibility levels at human NSC-specific DARs shown in Figure 3C. (C) Scatterplot depicting differential accessibility levels of human NSC-specific peaks (Figure 3C) in cortical organoids. (D) Heatmap displaying chromatin accessibility, centered at human NSC-specific DARs, grouped into RGC-specific, shared and IPC-specific based on Figure 4C. (E) Barplot depicting the overlap between the species-NSC specific and conserved NSC DARs, and significantly active enhancers in RGCs, IPCs, both or neither based on cell type-specific MPRA assay^44^. The NHP and Cons DARs are subsampled to match the human-specific peaks. M, crab-eating macaque; G, gorilla; C, chimpanzee; H, human; Cons, conserved. (F) Heatmap depicting the MPRA signal of the human NSC-specific DARs overlapping with enhancers active in RGCs, IPCs or both. (G) Contact maps and accessibility tracks for RGCs and IPCs at the *ZFHX4 locus*. The dashed rectangle indicates a human NSC-specific open region overlapping with an active enhancer in RGCs, which interacts with the *ZFHX4* promoter specifically in RGCs.

Next, we asked if human NSC DARs are also cell type-specific in human organoids. We found that these regions were much more accessible in RGCs compared to IPCs (Figure 4B–D), with the majority of them being RGC-specific (2073), a smaller proportion accessible in both cell types (shared – 701) or only in IPCs (197). These results suggest that many of the human NSC DARs we identified are also accessible in RGCs from cortical organoids.

To further validate if the human NSC-specific regions also include bona fide enhancers, we overlapped these regions with a massively parallel reporter assay (MPRA) we previously performed in cortical organoids^44^ (Figure 4A). This analysis revealed that 38.75% of the human NSC-specific regions for which we could extract MPRA information (n=80) had enhancer activity in the organoids, with the majority (71%) being NSC-specific (Figure 4E–F). These results were significantly higher compared to other NSC-specific DARs, such as chimpanzee (18.75%) and even conserved NSC-specific DARs (28.2%).

This is exemplified at the *ZFHX4* locus, which is expressed specifically in RGCs in the human fetal cortex^57^ and has been implicated in neurodevelopmental disorders, such as intellectual disability^58^. Here, we identified one human NSC-specific enhancer, which is accessible specifically in RGCs but not in IPCs and is engaged in a cell type-specific loop with the *ZFHX4* promoter (Figure 4G). Furthermore, this enhancer was highly active in RGCs but not in IPCs based on our MPRA assay (RGCs p<2.2e-16; IPCs p=n.s).

Overall, these results validate our human NSC-specific regions in an orthogonal system, such as cortical organoids. They further highlight the cell type specificity of these DARs with most of them being accessible and active in RGCs consistent with their profile in the 2D NSC system. By integrating them with MPRA and Hi-C data, we were also able to generate a set of high-confident putative enhancers that could be prioritized for functional interrogation of human-specific GRNs.

### Predicting the evolution of CRE activity from the DNA sequence

Changes in CRE activity across species can arise due to divergence in cis (e.g., mutations in the DNA sequence) or in trans (e.g., biochemical rates, TF abundance, metabolism)^59^. To address this, we used the deep learning model ChromBPNet^25^ to predict chromatin accessibility directly from the DNA sequence in each species and condition separately (Figure 5A).

**Figure 5.**
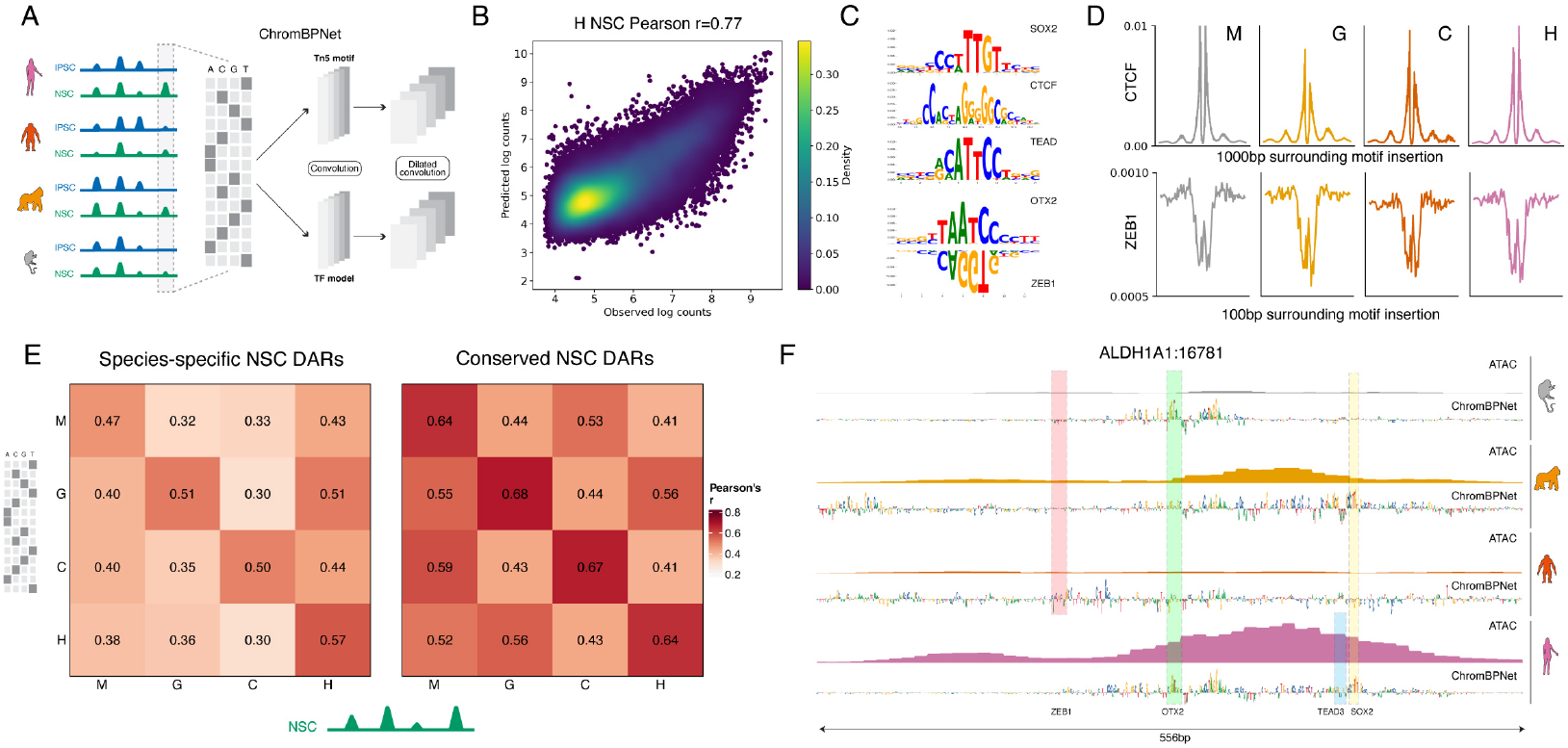
Deep learning model links changes in DNA sequence to evolution of the chromatin accessibility landscape. (A) Schematic of the deep learning model ChromBPNet^25^ used to predict chromatin accessibility from DNA sequence in each species and cell type separately. (B) Density scatter plot depicting the Pearson’s correlation between the measured and predicted log counts from human NSC peaks in held out chromosomes. (C) Top 5 TF-MODISCO contribution weight matrix (CMW) motifs derived from count contribution scores of the human NSC ATAC ChromBPNet no bias model. (D) Marginal footprint of the CTCF and ZEB1 motifs using ChromBPNet no bias predictions based on NSC ATAC in the corresponding species. (E) Cross-species model prediction for each class of species-specific or conserved NSC DARs. Shown is Pearson’s correlation between predictions of different models (y-axis) and measured accessibility in NSC of each species (x-axis). (F) ATAC-seq and nucleotide contribution score (ChromBPNet) tracks at the *ALDH1A1* putative CRE *locus*. The predicted motifs for ZEB1, OTX2, TEAD3 and SOX2 are highlighted in the respective species.

First, we evaluated the accuracy of the model by comparing the predictions to the true accessibility. We observed high correlations across all species and cell types (Figure 5B, S4A–B, median r=0.766), indicating that the model has learned the key underlying features associated with accessibility. Consistent with our previous observations, the main motifs predicted in NSC were SOX2, TEAD, OTX2 and ZEB1 (Figure 5C), while OCT4, NF-Y and ZIC were IPSC-specific (Figure S5C), which is in agreement with their described role in pluripotency^60–62^. The motifs in RGCs were comparable to the ones identified in human NSC but also included the TFs LHX2, NFI and RFX, while IPCs were associated with NEUROG2 and EOMES (Figure S5D), as previously described^44,51^.

Next, we asked if trans factors, such as the binding of TFs, are associated with differences in chromatin accessibility genome-wide. To address this, we performed motif footprinting for CTCF and ZEB1, embedding the corresponding motif in a randomized flanking sequence (Figure 5D). For both TFs we observed highly similar footprints using bias-corrected models in NSC across the four different species. This suggests that the binding of TFs (and the corresponding accessibility footprint) is comparable across primate evolution.

To address if DNA sequence alone is sufficient to explain the observed differences in chromatin accessibility, we compared the model predictions against the ground truth for species-specific NSC DARs versus conserved NSC DARs (based on the analysis in Figure 3C). Surprisingly, we observed that predictions for divergent DARs were less accurate than those for conserved DARs for the same species-model comparisons, regardless of which model was used (Figure 5E). Furthermore, the overall cross-model predictions (for example, the chimpanzee model asked to predict human accessibility given human sequence) were lower than those from the same model-species combinations. These results suggest that changes in DNA sequence alone are not sufficient to fully explain epigenome evolution.

To disentangle the complex interplay between changes in the linear genome and the epigenome context, we examined the nucleotide contribution scores at the *ALDH1A1* CRE *locus*, which is preferentially accessible in human NSC (Figure 3E). Consistent with our previous observations, we observed that the ZEB1 motif is present in all non-human primate (NHP) species but is absent in human, although this absence is only partially reflected in the importance scores (Figure 5F). Furthermore, we identified OTX2 and SOX2 motifs, which are conserved across all four species, while the TEAD motif was present only in human (Figure 5F). Therefore, such combination of repressor TF motif loss and gain of a novel activator TF motif could have contributed to the human-specific activity of this CRE.

Overall, these results suggest that changes in the DNA sequence are not sufficient to fully explain the epigenome evolution associated with chromatin accessibility across primate neurogenesis. Instead, complex regulatory grammar at CREs, the local context and potentially the proximity to other genomic elements contribute to species and cell type-specific differences.

### Dynamics of the 3D genome during primate neocortex development and evolution

In addition to chromatin accessibility, changes in global 3D genome organization or specific interactions between regulatory elements, such as enhancers and promoters, could have contributed to epigenome evolution in the context of primate brain development. Therefore, we profiled the 3D nuclear architecture using high-resolution Hi-C in both IPSC and NSC in each of the four species. We sequenced more than 6.29 billion reads in total, obtaining ∼513 million unique contacts per condition on average.

We found that global 3D genome organization was highly similar in IPSC across all four species across multiple levels examined: contact frequencies as a function of the genomic distance (Figure 6A – left), compartment strength (Figure S6A–B) and TADs (Figure S6C). However, we observed surprising divergence in 3D genome organization upon differentiation, specifically in chimpanzee and gorilla. Chimpanzee NSC were characterized by weaker mid-range and stronger long-range interactions (Figure 6A – right) and intermediate compartment strength (Figure 6B–C) but very strong insulation at TAD boundaries (Figure 6D, S6C). Conversely, gorilla NSC exhibited stronger compaction in the 100kb-1mb range, weaker long-range contacts as well as lower compartment score and chromatin insulation (Figure 6B–D, S6C). This was accompanied by decreased intra-TAD contacts and increased inter-TAD contacts, as well as fewer trans chromosomal interactions during differentiation, specifically in gorilla, which exhibited the opposite trend of all the other species (Figure 6D–E, S6D).

**Figure 6.**
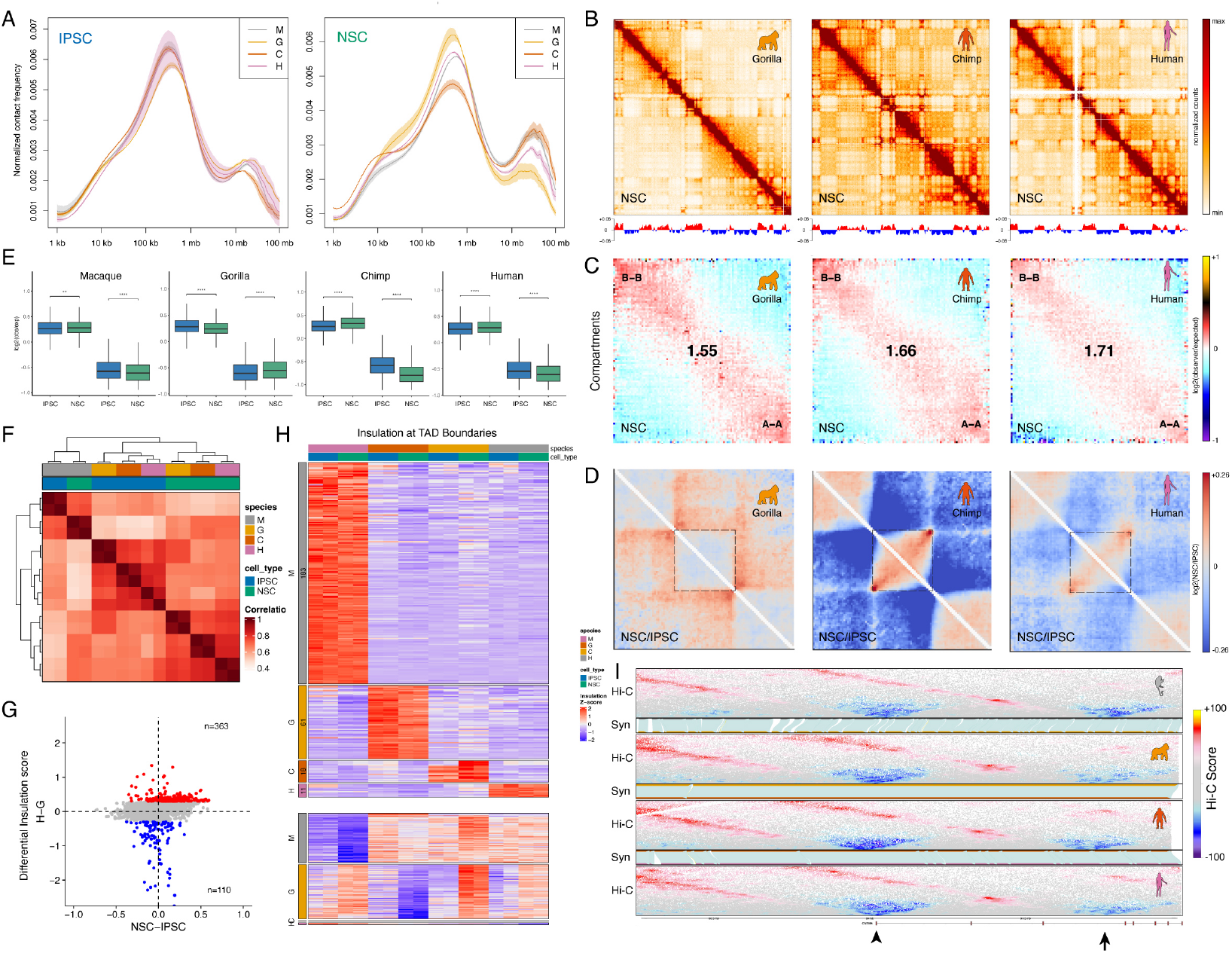
Dynamics of the 3D genome during primate brain evolution. (A) Normalized contact frequency in logarithmic bins (log2) for primate IPSC and NSC. Lines: means of biological replicates; semi-transparent bands: SEM. (B) Observed contact matrices for chr7 at 250kb resolution (top) and the first eigenvector at 100kb resolution (bottom) for gorilla, chimpanzee, and human NSC. (C) Saddle plots showing the average contact enrichment in NSC at pairs of 100kb *loci* arranged by their eigenvalue. Numbers represent the compartment strength. (D) Relative log ratio (NSC/IPSC) of the contact enrichment aggregated across TADs. (E) Quantification of the average intra- and inter-contact enrichment at TADs for primate IPSC and NSC. (F) Pairwise correlation matrix displaying Spearman’s correlation coefficient for chromatin insulation at the union of TAD boundaries across species. (G) Scatterplot showing a direct four-way comparison of changes in insulation across human vs gorilla and NSC vs IPSC. (H) Heatmap depicting groups of species-specific differentially insulated TAD boundaries. (I) Contact enrichment (Hi-C) in NSC and synteny maps (Syn) at the *CNTN5 locus* for all four primate species. The previously identified conserved TAD boundary is marked by an arrowhead, while the arrow on the right highlights the reported human-specific TAD boundary^21^. Note that the insulation appears highly conserved across both regions.

To determine the conservation of TADs across primate species, we used the insulation score^63^. We observed high reproducibility between replicates of the same sample and found that insulation across TAD boundaries was grouped mostly by cell type, with only macaque clustering separately (Figure 6F). Comparing human and gorilla, we observed that species-specific boundaries were very rare (473/9724 or 5%) and were not associated with a cell type bias (Figure 6G). Consistent with these results, when we expanded our analysis to include all species and cell types we also observed very few differentially insulated boundaries, suggesting a very strong conservation of TADs in primate evolution (Figure 6H).

To further clarify this, we examined the *CNTN5 locus*, which has been reported to contain a conserved and a human-specific TAD boundary^21^. However, our analysis indicated that both boundaries are conserved, and we did not observe any species-specific differences in chromatin insulation in either of them (Figure 6I, S6E), further highlighting the importance of considering multiple species and matched cell types when comparing the epigenetic landscape associated with evolution.

Overall, our analysis reveals novel dynamics of global chromatin organization during neural differentiation in primate evolution. Importantly, it leads to several novel conclusions: a) chimpanzee and especially gorilla 3D genome organization in NSC is different at the global scale compared to human and macaque; b) chromatin insulation dynamics in closely related species are associated primarily with cell type differences rather than the phylogenetic order and c) the vast majority of TAD boundaries appear conserved across species.

### Genome-wide comparative analysis of enhancer-promoter interactions

Having identified an evolutionary convergent and divergent set of putative CREs, we next asked if changes in enhancer-promoter (E-P) interactions also contribute to epigenome evolution. Similar to the analysis based on the insulation score, we observed high reproducibility between replicates of the same sample and found that E-P interactions were grouped mostly by cell type, with only macaque clustering separately (Figure 7A).

**Figure 7.**
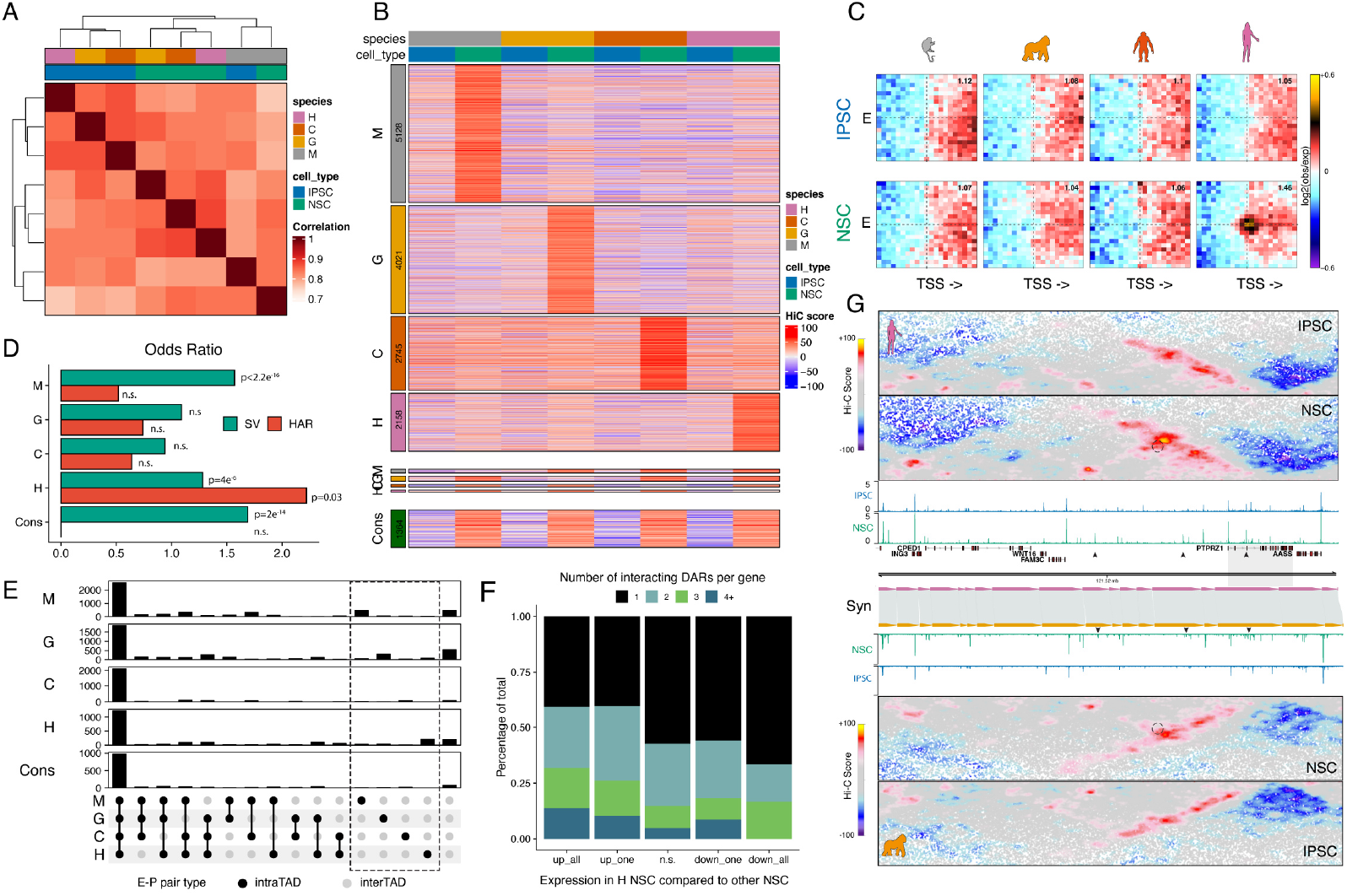
Comparative analysis of enhancer-promoter interactions. (A) Pairwise correlation matrix displaying Spearman’s correlation coefficient for pairs of distal peaks and promoters (E-P for short henceforth). (B) Heatmap depicting differential NSC E-P interactions. (C) Aggregate contact enrichment for human NSC-specific E-P pairs (n = 2158). Number in top-right corner indicates the ratio of the center enrichment to the mean of the four corners. E, enhancer. (D) Odds ratio of the different categories of E-Ps shown in B) overlapping with fhSVs or HARs. (E) Upset plot showing the intersection between different E-P classes and TADs in each species. The dashed rectangle indicates the interactions that are intra-TAD in one species but inter-TAD in all the other species. Note the increase of inter-TAD E-P interactions per group for each corresponding species. (F) Stacked barplot showing the number of interacting human NSC DARs per gene (based on Fig. 3H) that are downregulated in all species (down_all), downregulated in only one species (down_one), not significant (n.s.), upregulated in one (up_one) or in all species (up_all) when comparing expression in human versus NHP NSC. (G) Contact enrichment and synteny (Syn) plots for human and gorilla NSC at the *PTPRZ1 locus*. The dotted circle indicates a human NSC-specific E-P interaction identified in B) and associated with gene expression change. The small arrows point to the human NSC-specific putative enhancers. M, crab-eating macaque; G, gorilla; C, chimpanzee; H, human; Cons, conserved.

Next, we asked if E-P interactions are primarily conserved or species-specific. We identified 14052 NSC and 11416 IPSC species-specific E-P interactions (Figure 7B, S7A), validating the specificity of our approach using aggregate Hi-C enrichments (Figure 7C). Importantly, these differential interactions (DI) represented only 1.5% and 1.2% of all E-P contacts, respectively, and were significantly less than the number of DARs we identified (19.35% and 11.9% for NSC and IPSC, respectively). These results suggest that 3D regulatory interactions are much more conserved across evolution compared to changes in the linear epigenome.

To explore possible mechanisms underlying the species-specific E-P interactions, we examined the enrichment of HARs at each anchor or fhSV contained within the loop. We found that HARs were significantly enriched in human-specific NSC but not IPSC E-P interactions (Figure 7D, S7B), consistent with recent findings identifying the rewiring of regulatory interactions associated with HARs^23^. However, only a minor fraction of the DI overlapped with HARs (0.3%), suggesting that many regulatory contacts have changed independently.

Next, we asked if species-specific regulatory contacts remain within the same TAD (intra-TAD) or shifted between TADs (inter-TAD) during evolution, addressing the contribution of enhancer hijacking^23,64,65^ to 3D epigenome evolution. We observed that the majority of the differential E-P contacts were intra-TAD across all species in both NSC and IPSC (Figure 7E, S7C). However, we also observed that species-specific E-P contacts were disproportionally associated with TAD transitions (Figure 7E – dashed rectangle) – e.g., human NSC-specific interactions became intra-TAD in human but were inter-TAD in all other three species. To further explore enhancer hijacking, we examined the dynamics across human NSC E-P pairs based on the ABC model. We observed that the majority of these interactions (86%) were localized to the same TAD in all species (Figure S7D). These results suggest that although enhancer hijacking contributes to dynamic 3D regulatory interactions, the majority of changes occur across intra-TAD pairs.

Finally, we asked how changes in E-P interactions are linked to gene expression. Contrary to our analysis based on DARs (Figure 3H), we found that genes associated with human NSC-or IPSC-specific regulatory interactions were not differentially expressed across species (Figure S7E-F). However, we observed that most genes upregulated in human NSC compared to NSC from other species interacted with multiple DARs within the same TAD, while genes which were not changed or downregulated were primarily linked to a single DAR (Figure 7F). This result suggests that potential synergistic effects across divergent enhancers could be an important mechanism to explain species-specific gene regulation.

Finally, we identified several genes, such as *PTPRZ1*, a known marker of bRGCs^42,66^, where increased E-P interactions between its promoter and several human NSC-specific CRE correlated with increased expression (Figure 7H).

In conclusion, we developed a method to identify and quantitatively compare cell type-specific regulatory interactions across species. We identified thousands of species- and cell type-specific DI and identified HARs and enhancer hijacking as significant yet relatively minor contributions to these dynamics. Finally, we showed that changes in 3D regulatory interactions alone are not sufficient to explain gene expression dynamics. Instead, species-specific expression was associated with synergistic activation of multiple enhancers interacting in the 3D genome space, hinting towards a complex interplay between multiple regulatory modalities.

## Discussion

To understand the contribution of epigenome evolution to the expansion of the brain in primates, we comprehensively profiled transcriptome, chromatin accessibility, and 3D genome organization in IPSC and NSC from four primate species (crab-eating macaque, gorilla, chimpanzee and human). Using an integrative analysis, we identified both conserved and divergent gene regulatory networks that are associated with the evolution of the neocortex, validated the activity of human NSC-specific regulatory elements in cortical organoids, and used deep learning models to disentangle the relationship between DNA sequence and function.

Our comparative transcriptome analysis revealed that many genes from the WNT and NOTCH pathways are uniquely upregulated in human NSC, consistent with previous studies^19,34,39^. However, unlike human-specific genes such as *NOTCH2NL*, many of the genes we identified have NHP orthologs and are instead quantitively regulated. Additionally, we identified novel candidates with potential relevance, including *DMRT3* and *PTPRZ1*. Interestingly, Dmrt3 overexpression in mice enhances progenitor proliferation, leading to neocortical expansion^40^, suggesting a potential conserved mechanism. On the other hand, *PTPRZ1*, a known bRGC marker^42^, has been associated with the increased invasiveness of PTPRZ1-positive cells in glioblastoma^66^, indicating that it may be related to proliferation in both development and disease.

In contrast to previous studies, which have focused exclusively on gene expression, we also profiled chromatin accessibility and identified cell type- and species-specific CREs using a quantitative, reference-free approach. This revealed almost 7000 human NSC-specific CREs, significantly expanding the known regulatory elements associated with epigenome evolution in primates. We further validated these results by integrating them with RGC and IPC chromatin accessibility data from cortical organoids, validating the majority of them as RGC-specific. Finally, we compared the chromatin accessibility of these CREs with their ability to drive gene expression using a cell type MPRA in organoids^44^ and identified a set of high-confidence putative enhancers for further functional validation. Importantly, while changes in chromatin accessibility correlated overall with species-specific gene expression, many predicted target genes were not differentially expressed, likely due to compensatory^67,68^ effects.

Focusing on the molecular mechanism of epigenome evolution, we identified the gain of activator TF motifs alongside the loss of repressor TF motifs as one of the main predictors associated with species-specific changes. Consistent with previous studies, FOS::JUNB^44,49–51^ and TEAD^51,52^ motifs were enriched in NSC-specific DARs, supporting their role in NSC biology, while NR1H2::RXRA enrichment in human NSC DARs highlights the importance of retinoic acid in human neocortex specification^20,41^. In contrast, the ZEB1 repressor motif was uniquely lost in human NSC DARs. Notably, ZEB2, which shares a very similar motif to ZEB1, has been implicated in human brain evolution and has been linked to differences in the duration of the neuroepithelial transition^69^. Furthermore, we identified the TE family SVA as significantly enriched in human NSC DARs. SVAs represent the youngest group of retrotransposons in the human genome and have been previously linked to gene regulation in the context of pluripotency^70^. In human pluripotent stem cells, active SVAs are enriched in motifs for the TF YY1, which has been previously found to contribute to enhancer-promoter interactions and higher-order chromatin organization^71^, suggesting novel links between TE regulation and multimodal epigenome evolution in the context of human brain expansion.

In recent years, many studies have used deep learning models to explore the enhancer codes behind species differences^9,23,24,26^, although only a few used a multi-species-cell type context. Using the deep learning model ChromBPNet^25^, we observed that, although changes in accessibility are associated with differences in the underlying linear genome, this might not be sufficient to fully explain epigenome evolution. Instead, the differences likely arise from a complex regulatory grammar at CREs, influenced by local context and potential proximity to other genomic elements. Alternatively, epi-conserved regions might be robust to changes in sequence than more divergent sequences, leading to more accurate predictions for the conserved same-species model. In the future, exposing the model to multiple species and cell types and training a joint model incorporating different epigenetic modalities may increase the sensitivity, especially in closely related contexts.

Our multi-species, cell type-specific analysis revealed a unique pattern of global chromatin organization in gorilla cortical development, in which compartments and insulation at TAD boundaries become less pronounced upon differentiation. Although we do not yet know the exact reason for this interesting phenomenon, changes in the expression of the cohesin complex or loop extrusion dynamics might explain the differences observed. Furthermore, our comprehensive experimental setup confirmed the strong conservation of TAD boundaries across closely related primate species, as previously reported^72–75^. This observation contradicts recent findings, reporting a large number of human-specific TADs based on a pairwise comparison between macaque and human using micro-dissected fetal tissue^21^. This discrepancy underscores the importance of using quantitative models to measure differences in chromatin insulation^75^ instead of intersecting TAD boundary coordinates.

Finally, we introduced a novel approach to quantitatively compare E-P interactions across species. We found that regulatory interactions are much more dynamic across evolution than insulation at TAD boundaries but more conserved changes at the linear epigenome (e.g., chromatin accessibility). Furthermore, in contrast to E-P changes across cell types^51^, species-specific interactions were not generally accompanied by gene expression changes, again pointing to potential compensatory effects and highlighting the importance of multi-level molecular regulation. Importantly, we found that genes upregulated in human NSC interact with multiple differentially accessible regions (DARs) within the same TAD, whereas genes that are unchanged or downregulated are typically linked to a single DAR (Figure 7F). This observation implies that synergistic interactions among enhancers— where multiple regulatory elements cooperate to boost transcriptional output—may be a key mechanism driving species-specific gene regulation. Such enhancer synergy has been shown to confer robust gene expression in developmental contexts^67,68^ and our findings extend this concept to the evolution of the primate epigenome. These results underscore the potential importance of enhancer cooperation in shaping the complex gene regulatory networks that underlie human-specific neurogenic programs.

Recent studies examining HARs and 3D genome organization have reached conflicting conclusions about the conservation of the target genes and the importance of enhancer hijacking^23,64,65^. We observed that HARs are disproportionally enriched in the differential E-P interactions, yet they were only a very small minority of the overall differential interactions, pointing to other mechanisms involved in establishing species-specific 3D genome organization.

In summary, we established a new paradigm for quantitatively comparing the epigenome at multiple molecular levels, with the unique advantage of including multiple closely related species and ensuring cell type specificity. This lays the groundwork for the identification of novel human-specific molecular mechanisms and targets that can be further validated to deepen our understanding of the evolution of human traits. The extension of this methodology to single-cell comparative epigenomic studies using cortical organoids and fetal brain tissues, where available, will allow for studying cell types that are difficult to generate *in vitro*. This would also enable comparing evolutionary changes across different cell populations and developmental stages, significantly increasing the information we can extract and further advancing our understanding of brain evolution.

## Author Contributions and Notes

S.V. performed experiments. S.V., F.C. and B.B. analysed the data. T.D. established cross-species multimodal visualization. S.D. and W.E. provided the IPSC lines. B.B. supervised the project. S.V. and B.B. wrote the manuscript with input from all authors.

## Acknowledgments

Sequencing was performed at the Helmholtz Munich by the NGS-Core Facility and cell sorting was done at the Flow Cytometry Core Facility at the Biomedical Center, Ludwig-Maximilians-Universität. Work in the group of B.B. was supported by the Helmholtz Pioneer Campus, DFG priority program SPP2202 (BO 5516/1-1), ERA-NET Neuron (MOSAIC) and European Research Council Consolidator Grant (EpiCortex, 101044469). We thank Prof Magdalena Götz and Prof Anshul Kundaje for the useful discussions and input.

## Materials and Methods

### Cell culture and neural stem cells (NSC) generation

#### Human induced pluripotent stem cells (hIPSC)

The male hIPSC line (ISFi001-A) was provided by the IPSC Core Facility Helmholtz Munich^84^ and generated from human foreskin fibroblasts by transfection of five mRNA reprogramming factors (*Oct4, Sox2, Klf4, Lin28 and c-Myc*). Chromosomal analysis of the HMGU1 fixed cell suspension, performed by the Cytogenetics Laboratory of the Cell Guidance Systems (Cambridge, UK), revealed an apparently abnormal mosaic male karyotype. Five out of 20 cells examined contained an apparent duplication in the proximal long arm of one chromosome 20, band q11.2q11.2. However, this is a recurrent structural chromosomal abnormality reported in stem cell lines and has been associated with increased cell proliferation.

Cells were cultured at 37°C (5% CO_2_) on Matrigel (Corning, Cat. N.: 354277) coated plates (StemCell Technologies, Cat. N.: 38016) in mTeSR plus medium (StemCell Technologies, Cat. N.: 100-0276), and passaged as colonies using ReLeSR (StemCell Technologies, Cat. N.: 05872) or Gentle Cell Dissociation Reagent (StemCell Technologies, Cat. N.: 07174). Prior to each round of differentiation or before collection, hIPSC cultures were tested for mycoplasma contamination using LookOut Mycoplasma PCR Detection Kit (Sigma, Cat. N.: MP0035) and validated for pluripotency markers by immunohistochemical staining using the Human Pluripotent Stem Cell 3-Color Immunocytochemistry Kit (R×D Systems, Cat. N.: SC021).

#### Non-human primate induced pluripotent stem cells (IPSC)

The male chimpanzee IPSC line (WB01) was provided by Dr. Sebastian Diecke (MDC, Berlin) and generated from fibroblasts by Sendai virus-mediated reprogramming. The male gorilla IPSC line (55C1) derived from urinary stem cells and reprogrammed using Sendai virus was described and characterized before^28^. The female crab-eating macaque (= cynomolgus monkey = *Macaca fascicularis*) IPSC line (56A1) derived from fibroblasts and reprogrammed using Sendai virus had not been described before, but a clone from the same individual (56B1) has been described in the context of the generation of inducible KRAB-dCas9 from primates^85^. Chromosomal analyses of all the IPSC lines in a fixed cell suspension form were performed by the Cytogenetics Laboratory of the Cell Guidance Systems (Cambridge, UK). They have revealed the following: WB01 has an apparently normal male karyotype in all 20 cells examined; 55C1 has an apparently abnormal mosaic male karyotype because three of the 20 cells examined contained additional material of unidentified origin attached to the short arm of one chromosome 15 (breakpoint p11) and the remaining 17 metaphase cells examined had an apparently normal male karyotype; finally, 56A1 has shown an apparently normal female karyotype in all 20 cells examined.

Cells were cultured at 37°C (5% CO_2_) on Matrigel (Corning, Cat. N.: 354277) coated plates (StemCell Technologies, Cat. N.: 38016) in mTeSR plus medium (StemCell Technologies, Cat. N.: 100-0276). IPSC colonies were passaged using ReLeSR (StemCell Technologies, Cat. N.: 05872) or Gentle Cell Dissociation Reagent (StemCell Technologies, Cat. N.: 07174). Prior to each round of differentiation or before collection, IPSC cultures were tested for mycoplasma contamination using the LookOut Mycoplasma PCR Detection Kit (Sigma, Cat. N.: MP0035) and validated for pluripotency markers by immunohistochemical staining using the Human Pluripotent Stem Cell 3-Color Immunocytochemistry Kit (R&D Systems, Cat. N.: SC021).

#### Directed differentiation of primate IPSC to dorsal NSC

For the differentiation of NSC from primate IPSC, we followed a previously published protocol^29^ with some adaptations^30^. Briefly, IPSC were cultured at 37°C (5% CO_2_) on Matrigel (Corning, Cat. N.: 354277) coated 10cm dishes (Corning, Cat. N.: 353803) in mTeSR plus medium (StemCell Technologies, Cat. N.: 100-0276) to ∼100% confluence. At this point, the IPSC colonies were dissociated into single cells using pre-warmed StemPro Accutase Cell Dissociation Reagent (ThermoFisher, Cat. N.: A1110501) and resuspended in mTeSR plus medium (StemCell Technologies, Cat. N.: 100-0276) supplemented with 10µM of ROCK inhibitor Y-27632 (Sigma, Cat. N.: SCM075) to prevent cell death. Single cells were seeded at 1:1 ratio in a 10cm dish (Corning, Cat. N.: 353803), coated with growth factor-reduced Matrigel (Corning, Cat. N.: 354230) diluted 1:300 in DMEM/F-12 media (Gibco, Cat. N.: 11330-032) and incubated at 37°C (5% CO_2_). After 24-48 hours, when the culture covered the entire surface of the dish, the medium was replaced with neural induction medium containing a 1:1 mixture of DMEM/F-12 GlutaMAX (Gibco, Cat. N.: 31331-028) and Neurobasal medium (Gibco, Cat. N.: 21103-049) supplemented with 0.5% N2 Supplement (Life Technologies, Cat. N.: 17502-048), 0.025% Insulin (Sigma, Cat. N.: I9278), 1% B-27 Supplement (Life Technologies, Cat. N.: 17504044), 0.5% GlutaMAX Supplement (Life Technologies, Cat. N.: 35050-061), 0.5% MEM-NEAA (Life Technologies, Cat. N.: 1140-050), 1% Pen-strep (Gibco, Cat. N.: 15140122), and 0.1% b-mercaptoethanol (Gibco, Cat. N.: 31350010), with the addition of 1µM Dorsomorphin (StemCell Technologies, Cat. N.: 72102) and 10µM SB-431542 (StemCell Technologies, Cat. N.: 72232). Cells were cultured in these media until day 10, with daily media changes, when a uniform neuroepithelial sheet should be visible. On day 10, cells were dissociated with pre-warmed StemPro Accutase Cell Dissociation Reagent at seeded at a ratio of 1:3-1:5 on poly-L-ornithine/laminin-coated 10cm dishes – Poly-L-ornithine (1:500; Sigma, Cat. N.: p3655), Laminin (1:200; Sigma, Cat. N.: l2020) – and cultured for 24 hours in the same induction media as the previous 10 days with the addition of 10µM of ROCK inhibitor Y-27632. On days 11, 12 and 14, the media were replaced with fresh N3 media containing a 1:1 mixture of DMEM/F-12 GlutaMAX (Gibco, Cat. N.: 31331-028) and Neurobasal medium (Gibco, Cat. N.: 21103-049) supplemented with 0.5% N2 Supplement (Life Technologies, Cat. N.: 17502-048), 0.025% Insulin (Sigma, Cat. N.: I9278), 1% B-27 Supplement (Life Technologies, Cat. N.: 17504044), 0.5% GlutaMAX Supplement (Life Technologies, Cat. N.: 35050-061), 0.5% MEM-NEAA (Life Technologies, Cat. N.: 1140-050), 1% Pen-strep (Gibco, Cat. N.: 15140122), and 0.1% b-mercaptoethanol (Gibco, Cat. N.: 31350010). On day 15, NSC were dissociated using pre-warmed StemPro Accutase Cell Dissociation Reagent and either collected for the subsequent assay or seeded onto poly-L-ornithine/laminin-coated 10cm dishes at a ratio of 1:2.5-1:3 for longer culture (N3 media + 10µM of ROCK inhibitor Y-27632). After 24 hours in the same induction media as the previous 10 days with the addition of 10µM of ROCK inhibitor Y-27632. On day 16, the media was replaced with fresh N3 media and the cells were splitted every 6 days until day 31, as previously described.

#### Cortical organoids (COs) generation

For the generation of human COs, we used a protocol previously established in our laboratory^44^. Briefly, hIPSC were cultured on Matrigel (Corning, Cat. N.: 354277) coated 10cm dishes (Corning, Cat. N.: 353803) in mTeSR plus medium (StemCell Technologies, Cat. N.: 100-0276) at 37°C (5% CO_2_) until 80-90% confluence. The day before reaching the confluence, hIPSC were pre-treated with 1% DMSO (Sigma, Cat. N.: D2650) and the medium was switched to the complete Essential 8 Medium (Life Technologies, Cat. N.: A1517001). After 24 hours, single cells suspension was generated using Gentle Cell Dissociation Reagent (StemCell Technologies, Cat. N.: 07174) and 1×10^4^ single cells were seeded into AggreWell 800 plate (StemCell Technologies, Cat. N.: 34815), pre-treated with 500µl of the Anti-Adherence Rinsing Solution (StemCell Technologies, Cat. N.: 07010), in complete Essential 8 Medium with 10µM of ROCK inhibitor Y-27632 (Sigma, Cat. N.: SCM075) to promote survival. Cells were centrifuged at 100g for 3min at room temperature into the microwells and, after 24 hours, embryo bodies were harvested, transferred to ultra-low attachment 10cm dishes (Corning, Cat. N.: 3262) and cultured in Essential 6 Medium (Life Technologies, Cat. N.: A1516401) supplemented with 2.5µM Dorsomorphin (StemCell Technologies, Cat. N.: 72102), 10µM SB-431542 (StemCell Technologies, Cat. N.: 72232) and 2.5µM XAV-939 (Tocris, Cat. N.: 3748) (the latter only for 5 days). Medium was changed daily, except for day 1, and embryo bodies were embedded in a drop of Matrigel (Corning, Cat. N.: 354234) on day 7 and cultured in differentiation media (without vitamin A) containing a 1:1 mixture of DMEM/F-12 (Gibco, Cat. N.: 11330-032) and Neurobasal medium (Gibco, Cat. N.: 21103-049) supplemented with 0.5% N2 Supplement (Life Technologies, Cat. N.: 17502-048), 0.025% Insulin (Sigma, Cat. N.: I9278), 1% B-27 Supplement minus vitamin A (Life Technologies, Cat. N.: 12587010), 1% GlutaMAX Supplement (Life Technologies, Cat. N.: 35050-061), 0.5% MEM-NEAA (Life Technologies, Cat. N.: 1140-050), 1% Pen-strep (Gibco, Cat. N.: 15140122), and 0.1% b-mercaptoethanol (Gibco, Cat. N.: 31350010) for 4 days. The medium was then switched to differentiation medium (with vitamin A) containing a 1:1 mixture of DMEM/F-12 and Neurobasal medium supplemented with 0.5% N2 Supplement, 0.025% Insulin, 1% B-27 Supplement (Life Technologies, Cat. N.: 17504044), 1% GlutaMAX Supplement, 0.5% MEM-NEAA, 1% Pen-strep, and 0.1% b-mercaptoethanol, with media changes every 3-4 days, and the COs cultured on a shaker (neoLab, Cat. N.: 7-0950) at 85rpm. COs were harvested after 45 days and dissociated using the Papain-based Neural Tissue Dissociation Kit (Miltenyi Biotec, Cat. N.: 130-092-628) according to the manual dissociation protocol.

#### Immunofluorescence of NSC

For immunofluorescence, cells were differentiated directly on 13mm round coverslips (VWR, Cat. N.: 630-2118) using the same protocol as described in the previous section. Cells grown on 13mm round coverslips were washed in PBS, fixed in freshly prepared 4% formaldehyde (ThermoFisher, Cat. N.:28906) in PBS at room temperature for ∼10min and washed two or three times with PBS for 5-7min. After fixation, cells on coverslips were washed with 0.5% Triton X-100 (Sigma Aldrich, Cat. N.: X100) in PBS for 5min and incubated in PBS blocking buffer containing 10% horse serum (Sigma-Aldrich, Cat. N.: H0146) and 0.5% Triton X-100 (Sigma Aldrich, Cat. N.: X100) for 45min at room temperature. Staining was performed either for 2 hours at room temperature or overnight at 4°C in the dark with anti-PAX6 (1:100 dilution; Biolegend, Cat. N.: 901301), anti-EOMES (1:150 dilution; R&D Systems, Cat. N.: AF6166) and anti-TUBB3-AlexaFluor647 (1:100; BD Biosciences, Cat. N.: 560394) antibodies. All antibodies were diluted in PBS blocking buffer. Cells were washed three times with PBS for 5min followed by secondary staining with donkey anti-rabbit-A488 (Thermo Scientific, Cat. N.: A11015), donkey anti-sheep-A555 (Thermo Scientific, Cat. N.: A32794), donkey anti-mouse-A647 (Thermo Scientific, Cat. N.: A31571) and DAPI (Thermo Scientific, Cat. N.: D1306), all diluted in blocking buffer (1:500), for 45min at room temperature in the dark. The coverslips were washed three times with PBS for 5min before mounting onto Superfrost Plus adhesive microscope slides (ThermoFisher, Cat. N.: J1800AMNZ) using Fluoromount-G (Invitrogen, Cat. N.: 00-4958-02). All images were acquired using a Zeiss LSM 710 confocal microscope.

#### Cell fixation

1×10^6^ cells/mL of IPSC, D15 NSC and dissociated D45 COs were fixed with 1% formaldehyde (ThermoFisher, Cat. N.:28906) in PBS for 10min at room temperature with slow rotation and the reaction was quenched with 0.2M Glycine (ThermoFisher, Cat. N.: 15527-013) for 5min at room temperature with slow rotation. The cells were then spun down at 500g for 5min at 4°C and washed once with PBS containing 1% BSA (Sigma-Aldrich, Cat. N.: B6917) and 0.01% RNAsin plus RNase inhibitor (Promega, Cat. N.: N261A). Fixed IPSC were directly used for the subsequent assays, while fixed NSC and cells from COs were first subjected to fluorescence-activated cell sorting before use.

#### Fluorescence-activated cell sorting (FACS)

To isolate pure NSC and human radial glial cells (RGCs) and intermediate progenitor cells (IPCs) from COs, we followed a protocol previously described^44,51^. Briefly, dissociated and fixed cells were first incubated in a permeabilization buffer consisting of 0.1% freshly prepared Saponin (Sigma-Aldrich, Cat. N.: SAE0073), 0.2% BSA (Sigma-Aldrich, Cat. N.: B6917), and 0.01% RNAsin plus RNase inhibitor (Promega, Cat. N.: N261A) in PBS for 15min at 4°C. The permeabilization buffer was removed by centrifugation at 2500g for 5min at 4°C, followed by staining against PAX6-AlexaFluor488 (1:40; BD Bioscience, Cat. N.: 561664) for NSC, and SOX2-PE (1:20; BD Biosciences, Cat. N.: 562195), PAX6-AlexaFluor488 (1:40; BD Bioscience, Cat. N.: 561664) and EOMES-eFluor660 (1:20; BD Bioscience, Cat. N.: 566749) for RGCs and IPCs, in staining buffer consisting of 0.1% freshly prepared Saponin, 1% BSA and 0.04% RNAsin plus RNase inhibitor in PBS for 1 hour at 4°C under slow rotation. Cells were washed twice for 5min in PBS containing 1% BSA and 0.01% RNAsin plus RNase inhibitor and then incubated for 10min at 4°C with slow rotation in PBS containing 0.5% BSA and 0.04% RNAsin plus RNase inhibitor, and DAPI (1:1000; ThermoFischer, Cat. N.: D1306) in the case of CO sample. For the COs sorting strategy, singlets were selected using forward and side scatter, followed by the identification of cells in G_0_-G_1_ by genomic content based on DAPI staining. These cells were then divided into SOX2^-^/EOMES^+^ for IPCs and SOX2^+^/EOMES^-^ for further subselection to SOX2^+^/PAX6^+^ for RGCs. In the case of NSC, singlets were selected on the gate identified by forward and side scatter and these cells were identified as NSC based on PAX6 expression (PAX6^+^). Cell sorting was performed on a FACSAria III (BD Biosciences; lasers: 405nm, 488nm, 561nm, 633nm) using a 100µm nozzle. After sorting, ∼10^5^ cells were either directly used for ATAC-seq, flash-frozen for Hi-C or RNA was extracted using the Quick-RNA FFPE kit (Zymo Research, Cat. N.: R1008) with Zymo-Spin IC Columns (Zymo Research, Cat. N.: C1004-250). FACS plots were generated using FlowJo (Version 10.9.0).

### IPSC, NSC, RGCs and IPCs multimodal profiling

#### RNA-seq

To generate RNA-seq libraries, RNA from ∼10^5^ fixed IPSC or fixed-sorted NSC was isolated using the Quick-RNA FFPE kit (Zymo Research, Cat. N.: R1008) in combination with Zymo-Spin IC Columns (Zymo Research, Cat. N.: C1004-250) according to the manual instructions starting from the tissue dissociation step. A DNase I treatment (Zymo Research, Cat. N.: E1010) was performed at the end to remove genomic DNA contamination, followed by RNA purification using the RNA Clean & Concentrator-5 kit (Zymo Research, Cat. N.: R1013) according to the instructions. The yield was quantified using the Qubit RNA HS Assay Kit (ThermoFisher, Cat. N.: Q32852) and a high RNA quality (RIN>7.5) was verified using the Bioanalyzer High Sensitivity RNA 6000 Pico Kit (Agilent, Cat. N.: 5067-1513).

Approximately 100ng of RNA was used to generate RNA libraries for all samples using the NEBNext® Single Cell/Low Input RNA Library Prep Kit (New England Biolabs, Cat. N.: E6420) according to the manual guidelines.

#### ATAC-seq

ATAC-seq was performed following the Omni-ATAC-seq protocol^46^, with some optimizations for the success of the protocol with fixed cells and minor modifications in the PCR amplification step and in the size selection of the resulting libraries. Briefly, 10^5^ fixed IPSC or fixed-sorted NSC, RGCs or IPCs (highly viable before fixation) were spun down at 500g for 5min at 4°C and resuspended by pipetting up and down three times in 100µl of cold ATAC Resuspension Buffer consisting of 10mM Tris-HCl pH 7.4 (ThermoFisher, Cat. N.: 15567027), 10mM NaCl (ThermoFisher, Cat. N.: AM9760G), 5mM MgCl_2_ (ThermoFisher, Cat. N.: AM9530G), 0.1% IGEPAL CA630 (Sigma-Aldrich, Cat. N.: I3021), 0.1% TWEEN^®^ 20 (Sigma-Aldrich, Cat. N.: P9416), 0.01% Digitonin (Promega, Cat. N.: G9441) in Nuclease-free water (ThermoFisher, Cat. N.: AM9937). Cells were incubated for 3min on ice, if necessary, the lysis time can be extended to 5 or maximum 10min, and the concentration and the viability were determined (viability should be approximately 0%). 5 x 10^4^ cells were transferred to a new tube and washed with 1ml of cold ATAC Wash Buffer containing 10mM Tris-HCl pH 7.4 (ThermoFisher, Cat. N.: 15567027), 10mM NaCl (ThermoFisher, Cat. N.: AM9760G), 5mM MgCl_2_ (ThermoFisher, Cat. N.: AM9530G), 0.1% TWEEN^®^ 20 (Sigma-Aldrich, Cat. N.: P9416) in Nuclease-free water (ThermoFisher, Cat. N.: AM9937). After centrifugation at 500g for 10min at 4°C, the supernatant was carefully aspirated, the cell pellet was resuspended by pipetting up and down six times in 50µl of Transposition mix consisting of 50% 2X TD Buffer (Illumina Tagment DNA Enzyme and Buffer Small Kit; Illumina, Cat. N.: 20034197), 5% TDE1 Tagment DNA Enzyme (Illumina Tagment DNA Enzyme and Buffer Small Kit; Illumina, Cat. N.: 20034197), 33% PBS, 0.01% Digitonin (Promega, Cat. N.: G9441), TWEEN^®^ 20 (Sigma-Aldrich, Cat. N.: P9416) in Nuclease-free water (ThermoFisher, Cat. N.: AM9937), and the reaction was performed at 37°C for 1 hour in a thermomixer at 1000rpm. After the transposition step, 50µl of 2X Reverse-crosslinking solution containing 100mM Tris-HCl pH 7.4, 2mM EDTA (Sigma-Aldrich, Cat. N.: E7889), 2% SDS (ThermoFisher, Cat. N.: AM9823), 0.4M NaCl, 1% proteinase K (New England Biolabs, Cat. N.: P8107S) in Nuclease-free water was added to the previous reaction and all was incubated overnight at 65°C in a thermomixer with 1000rpm. The reaction was purified using the DNA Clean & Concentrator-5 kit (Zymo Research, Cat. N.: D4014) according to the instructions with a 1:5 ratio of the purified DNA. The product was mixed with NEBNext Ultra II Q5 Master mix (New England Biolabs, Cat. N.: M0544L) and 1.2µM of each PCR primer^46^ in the PCR amplification reaction using the following program: 98°C for 30sec; (98°C 10sec, 65°C 90sec) x 12; 65°C for 5min; hold at 10°C. After the amplification, the library was purified twice using 0.55x and 1.25x AMPure XP beads (Beckman Coulter, Cat. N.: A63881) to obtain an average fragment size between 200bp and 1000bp.

#### Hi-C

For *in situ* Hi-C, we adapted the current protocol^74^. Briefly, frozen pellets of 10^5^ fixed IPSC or fixed-sorted NSC, RGCs or IPCs were thawed on ice and then lysed in 300µl of cold Hi-C Lysis Buffer consisting of 10mM Tris-HCl pH 8.0 (ThermoFisher, Cat. N.: 15568025), 10mM NaCl (ThermoFisher, Cat. N.: AM9760G), 0.2% IGEPAL CA630 (Sigma-Aldrich, Cat. N.: I3021), 1% proteinase inhibitors (Roche, Cat. N.: 11873580001) in Nuclease-free water (ThermoFisher, Cat. N.: AM9937). After incubation for 30min on ice, cells were spun down at 2500g for 5min at 4°C and the pellet was washed once with 500µl of cold Hi-C Lysis Buffer (2500g for 5min). Cells were permeabilized with 50µl of 0.5% SDS (Invitrogen, Cat. N.: AM9823) at 62°C for 10min, followed by an incubation at 37°C for 15min at 600rpm, with 145µl of Nuclease-free water and 25µl of 10% Triton X-100 to quench the SDS. Chromatin was digested with 500U DpnII enzyme (New England Biolabs, Cat. N.: R0176) and 25µl of 10X DpnII buffer at 37°C overnight at 600rpm. The reaction was incubated at 62°C for 20min, allowed to cool to room temperature and the sticky ends were filled with biotin by adding and mixing a master mix containing 37.5µl of 0.4mM biotin-14-dATP (Life Technologies, Cat. N.: 19524016), 1.5µl each of 10mM dCTP, 10mM dGTP and 10mM dTTP, and 8µl 5U/µl DNA Polymerase I, Large (Klenow) Fragment (New England Biolabs, Cat. N.: M0210). After incubation at 37°C for 1.5 hours with mixing, 900µl of ligation master mix was added (669µl Nuclease-free water, 120µl 10X T4 DNA ligase buffer (New England Biolabs, Cat. N.: B0202), 100µl Triton X-100, 6µl 20mg/ml BSA (New England Biolabs, Cat. N.: B9000) and 5µl 400U/µl T4 DNA ligase (New England Biolabs, Cat. N.: M0202)) and the reaction was incubated at 16°C for at least 4 hours with slow mixing. Proteins were digested with 20µl 800U/ml proteinase K (New England Biolabs, Cat. N.: P8107) and 120µl 10% SDS at 55°C for 30min at 600rpm, followed by reverse cross-linking at 68°C overnight with 130µl 5M NaCl. DNA was then purified by ethanol precipitation and sheared to ∼550bp DNA fragments using a Covaris S220 sonicator. Biotin pulldown was performed by incubating the sheared DNA with 100µl of 10mg/ml MyOne Streptavidin T1 beads (Thermo Fisher, Cat. N.: 65602) for 30min at room temperature with rotation, followed by removal of biotin from the beads and end repair by incubation for 30min at room temperature in a reaction mix consisting of 88µl 1X T4 DNA ligase buffer, 2µl 25mM dNTP mix, 5µl 10U/µl T4 Polynucleotide Kinase (New England Biolabs, Cat. N.: M0201), 4µl 3U/µl DNA Polymerase I (New England Biolabs, Cat. N.: M0203) and 1µl 5U/µl DNA Polymerase I, Large (Klenow) Fragment. Subsequently, A-tailing was performed using 90µl 1X NEBuffer 2 (New England Biolabs, Cat. N.: B7002), 5µl 10mM dATP and 5µl 5U/µl NEB Klenow Fragment exo-minus (New England Biolabs, Cat. N.: M0212) was performed at 37°C for 30min with mixing, followed by ligation of 2µl NextFlex index adapters (Bioo Scientific, Cat. N.: NOVA-514102) using 50µl 1X NEB Quick ligation reaction buffer (New England Biolabs, Cat. N.: B2200) and 2µl NEB DNA Quick ligase (New England Biolabs, Cat. N.: M2200) for 15min at room temperature with mixing. Between each of the incubation step, the samples bound to the streptavidin beads were washed twice with wash buffer containing 5mM Tris-HCl pH 7.4, 0.5mM EDTA, 1M NaCl, and 0.05% TWEEN^®^ 20 in Nuclease-free water, and once with the buffer of the subsequent reaction. Libraries were amplified directly on the streptavidin beads (in 47µl of 10mM Tris-HCl pH 8.0) using 55µl NEBNext Ultra Q5 II Master mix and 5µl of NEBNext Universal PCR Primer for Illumina (index primer sequence: 5’-AAT GAT ACG GCG ACC ACC GAG ATC TAC ACT CTT TCC CTA CAC GAC GCT CTT CCG ATC-s-T-3’) using the following program: 98°C for 30s; (98°C 10s, 65°C 75s) x 10; 65°C for 5min; hold at 10°C. After the amplification, the streptavidin beads were discarded, and the supernatant was purified twice using 0.7x AMPure XP beads to obtain an average fragment size of approximately 500bp.

#### Library QC and Sequencing

Library quantification was performed by qPCR using the NEBNext® Library Quant kit (New England Biolabs, Cat. N.: E7630). The library size distribution was assessed with the Agilent High Sensitivity DNA Kit (Agilent, Cat. N.: 5067-4626) on the Agilent 2100 Bioanalyzer. Sequencing was conducted on NovaSeq6000. Sequencing and QC metrics are listed in Supplementary Data Table 1.

## QUANTIFICATION AND STATISTICAL ANALYSIS

### Comparative transcriptome analysis

RNA-seq libraries were mapped and deduplicated using STAR^78^ with default settings. Either hg38 was used as the reference genome for all the species, or macFas6, gorGor6, panTro6 and hg38 were used as the reference genomes for crab-eating macaque, gorilla, chimpanzee and human, respectively. In the case of individual genome mapping, orthologous genes were identified based on the corresponding annotated gene name. DESeq2^79^ was used to determine differentially expressed genes (FDR < 0.1, log2fc = 1) based on species (crab-eating macaque, gorilla, chimpanzee and human) and cell type (IPSC and NSC). Genes specifically enriched or reduced in IPSC or NSC in a given species and differentially expressed in all the other species with a log2 fold change of at least 1 were then identified, together with the conserved ones. Genes enriched in IPSC and NSC in each of the species are listed in Supplementary Data Table 2. Functional enrichment analysis was performed using Cluster Profiler and visualized with enrichPlot^86^. Enrichment analysis was performed to assess the association between the cell type-specific genes upregulated in each of the species and previously identified human-specific structural variants (fhSVs)^45^ or high-confidence human accelerated regions (HARs)^23^ within a 5kb window. The odds ratio and statistical significance of the enrichment were calculated using Fisher’s exact test.

### Comparative accessibility analysis

ATAC-seq libraries were mapped and processed using the ENCODE pipeline^80^ with default settings. MacFas6, gorGor6, panTro6 and hg38 were used as the reference genomes for crab-eating macaque, gorilla, chimpanzee and human, respectively. Putative orthologous regulatory regions were identified using HALPER^48^ after generating reference-free Cactus multiple sequence alignments of the macFas6, gorGor6, panTro6 and hg38 genomes^47^. The orthologous regions were merged to generate a common peak set (248465 peaks) with the genomic coordinates of each corresponding species. DESeq2^79^ was used to calculate differentially accessible regions based on species (crab-eating macaque, gorilla, chimpanzee and human) and cell type (IPSC and NSC). Regions specifically open or close in IPSC or NSC in a given species and differentially accessible in all the other species with a log2 fold change of at least 1 were then identified, together with the conserved ones. Regions open in IPSC and NSC in each of the species are listed in Supplementary Data Table 2.

PhastCons30way conservation scores were used to extract sequence conservation of the NSC species-specific accessible and conserved regions.

Motif-based enrichment analysis was performed using the JASPAR2020 core vertebrate database. Transcription factor (TF) motif enrichment for TFs expressed in NSC or IPSC (FPKM>=1) was calculated using the monaLisa package^87^ using binomial test with 0.1 FDR cutoff and 0.2 enrichment cutoff.

Enrichment analysis was performed to assess the association between the cell type-specific open regions in each of the species and previously identified human-specific structural variants (fhSVs)^45^ or high-confidence human accelerated regions (HARs)^23^ within a 5kb window. The odds ratio and statistical significance of the enrichment were calculated using Fisher’s exact test.

The overlap between transposable elements and human NSC or conserved NSC-specific accessible regions was calculated using a two-sided Fischer’s exact test.

Human NSC Hi-C data were used to generate an Activity-by-Contact (ABC) model^56^, which was then applied to link human-specific accessible regions to genes.

### ChromBPNet

The deep learning DNA sequence model ChromBPNet^25^ was used to predict chromatin accessibility from the DNA sequence in each species and condition with the following details. Preprocessing involved merging two replicates to yield consolidated bam files, which were used as input to the models. Conserved peaks were called previously from the ENCODE pipeline. The reference genome fasta files and chromosome sizes from macFas6, gorGor6, panTro6 and hg38 were used for crab-eating macaque, gorilla, chimpanzee and human, respectively. For each species, we subsequently defined the following splits (fold_0) – chromosome 1, 3, 6 for testing, chromosome 8 and 20 for validation and the rest of the chromosomes for training. Background regions were defined from GC-matched peaks. Training was performed in two separate steps to generate species- and cell type-specific bias-factorized ChromBPNet models – resulting in a total of eight models. First, a bias model was trained using the default bias threshold factor of 0.5, and its performance was verified by ensuring that it captured the expected Tn5 bias motif. Next, the trained bias model was incorporated into the ChromBPNet training process. In this second step, a bias-corrected ChromBPNet model was trained. This was subsequently used to predict chromatin accessibility profiles of regions classified as conserved and non-conserved differentially accessible regions (DARs). *De novo* motif discovery was then performed using TF-MoDISco Lite to extract the top five motifs learned by the model. Additionally, *in silico* addition of motif profiles for CTCF and ZEB1 was carried out to understand, if any, the effect of specific TF binding on chromatin accessibility profiles.

### Hi-C data processing

Hi-C libraries were mapped and processed using Juicer^81^ and macFas6, gorGor6, panTro6 and hg38 were used as the reference genomes for crab-eating macaque, gorilla, chimpanzee and human, respectively. Normalization was performed using the Shaman package (https://tanaylab.bitbucket.io/shaman/index.html). Hi-C scores were calculated using a kNN strategy on combined replicates as described previously^63^ with a kNN parameter set to 100.

### Contact probability, compartments and compartment strength

We calculated and visualized the contact frequency as a function of genomic distance as described previously^63^. The dominant eigenvector of the contact matrices (250kb bins) was computed as described^72^ using scripts available at https://github.com/dekkerlab/cworld-dekker/. This was used to assign respective compartment identity (A or B compartments). The strength of interactions between A and B compartments was further interrogated by binning and ranking the log2 ratio of observed versus expected contacts. A ratio was calculated both between bins of the same (A-A, B-B) or different types (A-B), as detailed in a previous publication^63^. It represents the ratio between the sum of observed contacts within compartments A and B and the sum of contacts between compartments (AA+BB)/(AB+BA).

### Insulation, TAD boundary calling and average TAD contact enrichment

To define insulation based on observed contacts, we used the insulation score^63^, which was calculated on a combined contact map at 1kb resolution within a ±250kb region and then multiplied by (-1). TAD boundaries were identified as local maxima at 2kb intervals within regions where the insulation score was above the 90% quantile of the genome-wide distribution. To calculate insulation and contact enrichment within TADs, their coordinates were extended upstream and downstream by the TAD length and this distance was divided into 100 equal bins. The observed versus expected enrichment ratio was calculated in each resulting 100 × 100 grid (per TAD) and the average enrichment per bin was plotted. Cooltools^88^ was used for the average TAD representation.

### TAD conservation

TAD conservation was calculated using the insulation score to quantitatively measure the TAD boundaries as described previously. Putative orthologous TAD boundaries were identified using a similar approach to that used for chromatin accessibility^47,48^ and genomic regions with low mappability were removed. The orthologous regions were merged to generate a common set (10800 regions) with the genomic coordinates of each corresponding species. Changes in insulation between IPSC and NSC in a given species that were additionally altered in all the other species were identified. Regions insulated in NSC in each of the species are listed in Supplementary Data Table 2.

### Comparative regulatory interactions analysis

All possible E-P pairs within a ±500kb window were generated, and the interaction frequency between them (denoted as Hi-C score) in each species was calculated as described previously. Putative orthologous pairs were identified using HALPER^48^ after generating reference-free Cactus multiple sequence alignments of the macFas6, gorGor6, panTro6 and hg38 genomes^47^. The orthologous regions were merged to generate a common set (2060701 pairs) with the genomic coordinates of each corresponding species. Regulatory interactions, specifically strong or weak in IPSC or NSC in a given species and with a differential strength in all the other species, were then identified, together with the conserved ones. Strong interactions in IPSC and NSC in each of the species are listed in Supplementary Data Table 2. To validate the specificity of these enhancer-promoter pairs, observed and expected contacts pooled from replicates were extracted within an 80*80kb window. A higher log2(obs/exp) ratio represents stronger contact enrichment.

### Cross-species multimodal visualization

For the cross-species multimodal visualization, the human genomic coordinates of interest were converted into the orthologous genomic coordinates of the other primate species using halLiftover^82^. Per species and modality, the data values were extracted for those coordinates and plotted with ggplot with the plotting range adjusted to the maximum coordinate range across all species. The syntenies were plotted using an adjusted version of the plotMiro function from SVbyEye^83^.

