## Supplemental Figures for "3D Epigenome Evolution Underlies Divergent Gene Regulatory Programs in Primate Neural Development"

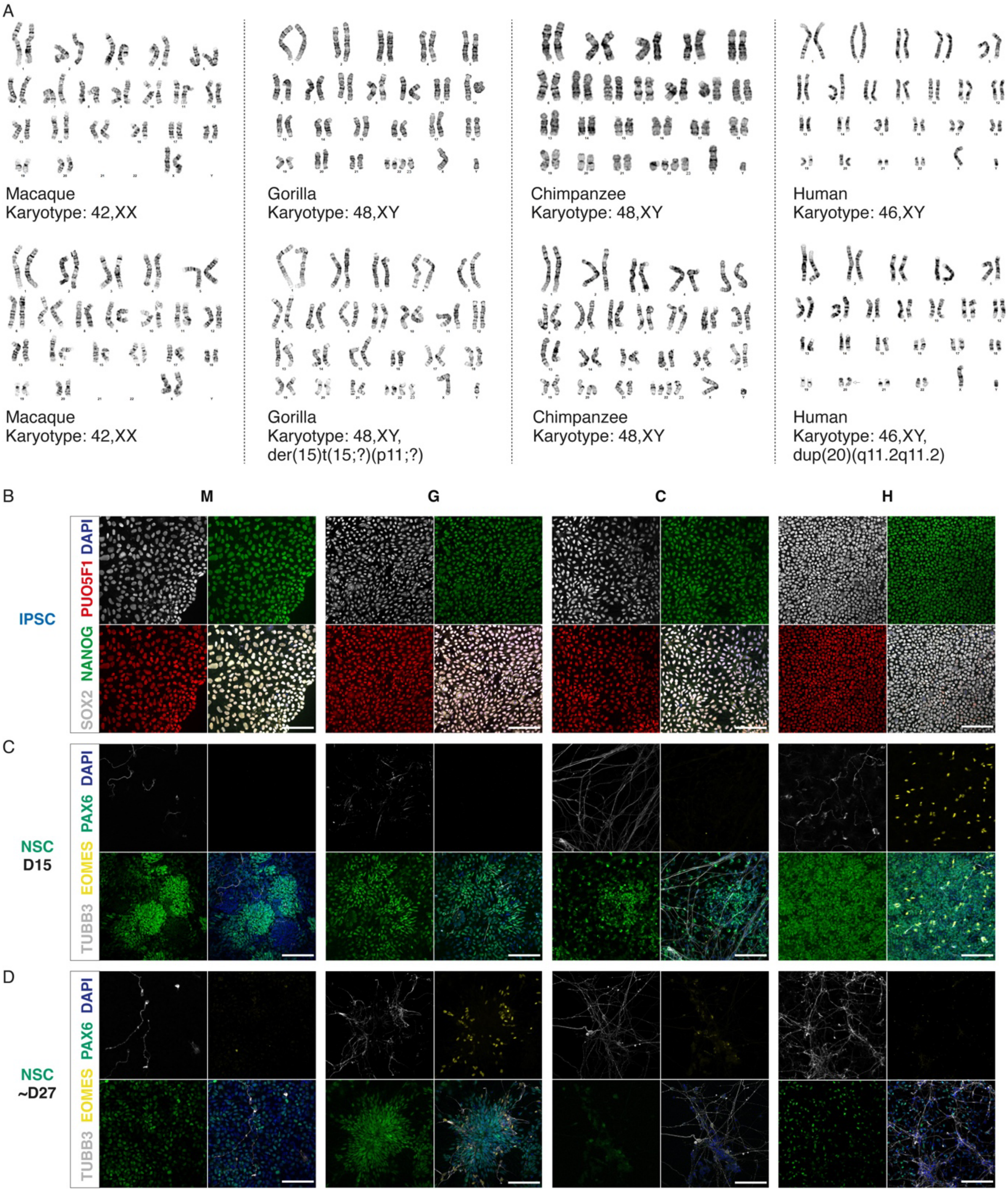


**Figure S1. Primate IPSC-derived neural stem cells as a model system to study human brain evolution**

(A) Two representative images of the IPSC karyotype for each of the four primate species. (B-D) Representative immunofluorescence images of primate IPSC (B) and primate NSC at day 15 (C) and at ~day 27 (D). M, crab-eating macaque; G, gorilla; C, chimpanzee; H, human. Scale bar, 100 µm.

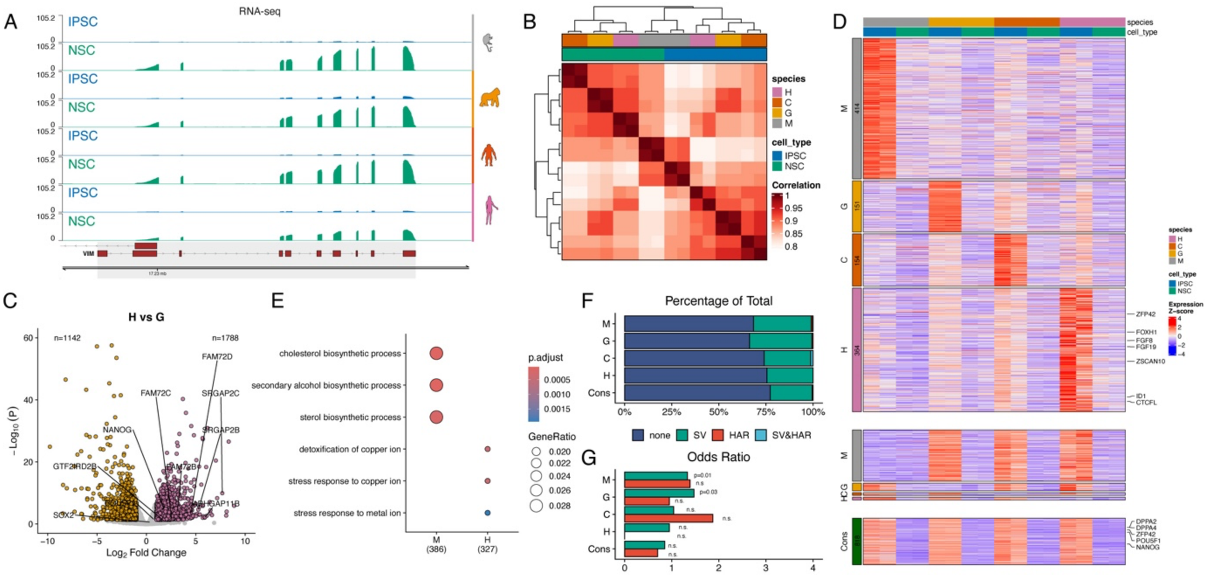


**Figure S2. Comparative transcriptomics reveals cell type- and species-specific gene expression**

(A) RNA-seq tracks of primate IPSC and NSC at the *VIM locus*. (B) Pairwise correlation matrix displaying Spearman's correlation coefficient for gene expression (reads mapped to the corresponding genome). (C) Scatterplot depicting differentially expressed genes (DEGs; false discovery rate (FDR) < 0.05) in human vs gorilla IPSC. (D) Heatmap depicting significantly upregulated (top), downregulated (middle) and conserved (bottom) primate IPSC genes. (E) Dot plot showing the gene ratios of the enriched gene ontology across the macaque and human upregulated genes in D). The color of the circles represents the Benjamini-Hochberg adjusted P value. (F-G) Percentage and odds ratio of upregulated or conserved genes in D that overlap with fhSVs[45] and/or zooHARs[23]. M, crab-eating macaque; G, gorilla; C, chimpanzee; H, human; Cons, conserved.

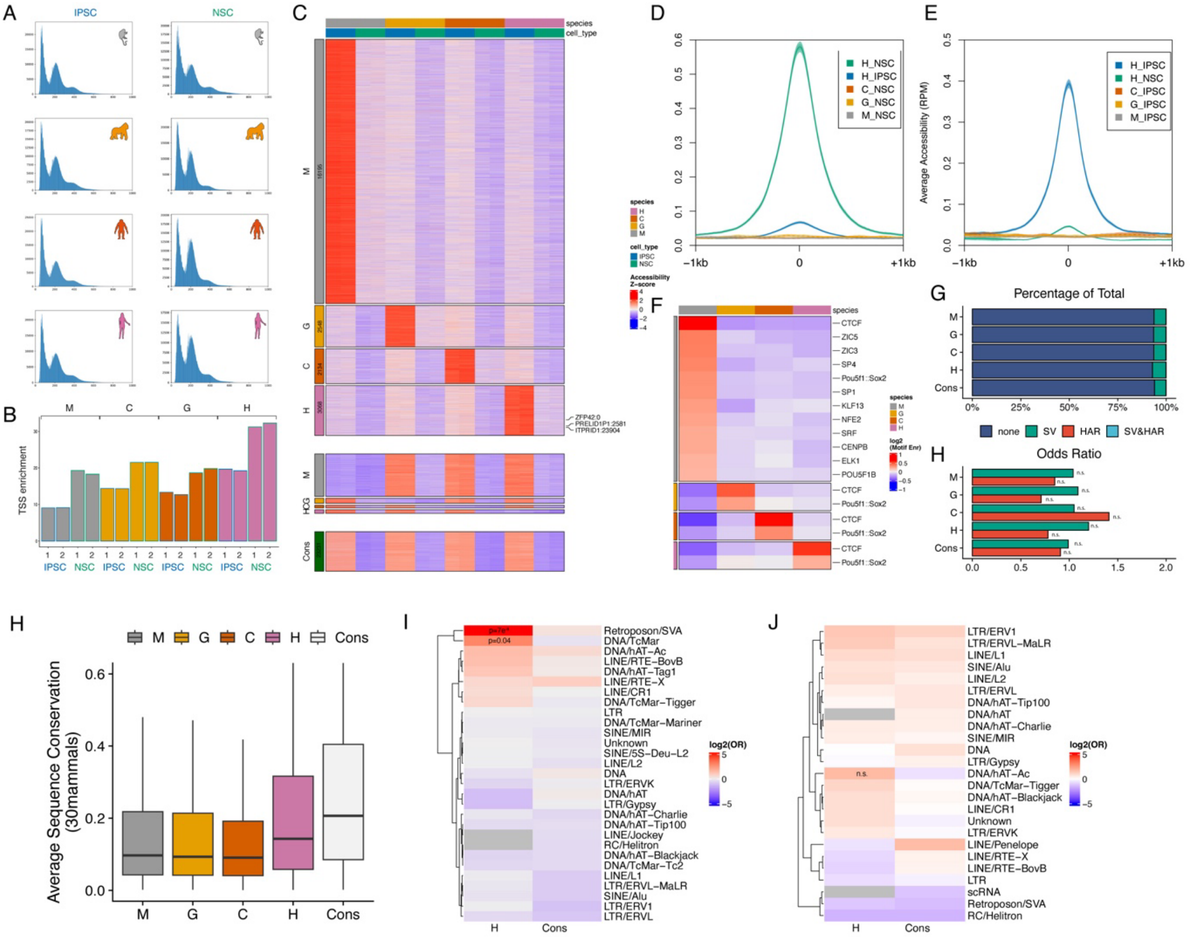

**Figure S3. Changes in TF motifs and TE underlie dynamic chromatin accessibility in primate evolution** (A) Distinct fragment length enrichments of the ATAC-seq libraries from primate IPSC and NSC. (B) Bar plot of the TSS enrichment of the ATAC-seq libraries from primate IPSC and NSC in two replicates. (C) Heatmap depicting differentially accessible regions (DARs) in primate IPSC. (D) Average accessibility levels at human NSC-specific peaks shown in 3D) for human IPSC, and primate NSC. (E) Average accessibility levels at human IPSC-specific peaks shown in 3D) for human NSC and primate IPSC. (F) Heatmap showing TF motif enrichment in the IPSC-species specific DARs regions in each species. (G-H) Percentage and odds ratio of IPSC DARs or conserved regions that overlap with fhSVs[45] and/or zooHARs[23]. (I) Average sequence conservation (phastcons30mammals) between different classes of DARs. (J-K) Odds ratio for the overlap between different classes of transposable elements and either human-specific or conserved DARs in NSC (J) or IPSC (K). M, crab-eating macaque; G, gorilla; C, chimpanzee; H, human; Cons, conserved.

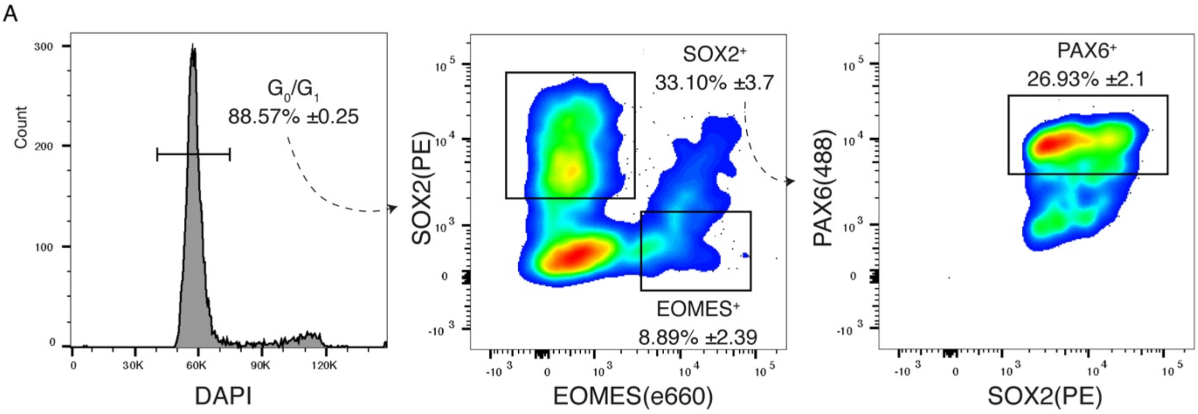


**Figure S4. Additional validation of the human-specific NSC DARs in human cortical organoids**

(A) Gating strategy used to isolate RGCs ($PAX6^+/SOX2^+/Eomes^-$) and IPCs ($SOX2^-/Eomes^+$) from D45 cortical organoids.

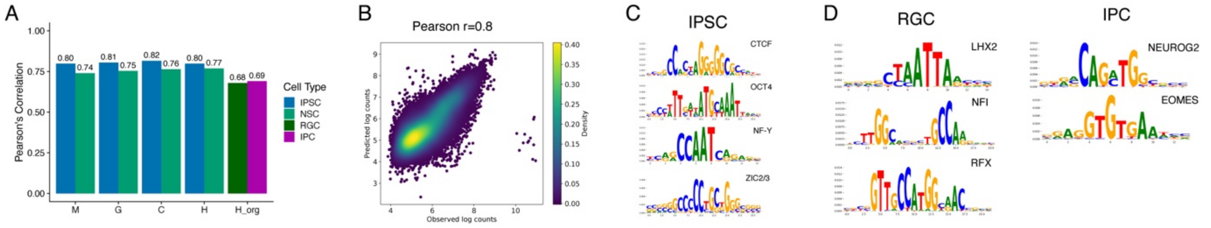


**Figure S5. Additional validation and predictions of the deep learning model**

(A) Barplot depicting the Pearson's correlations between the measured and predicted log counts in held out chromosomes in each species cell type combination. (B) Density scatter plot depicting the Pearson's correlation between the measured and predicted log counts from human IPSC in held out chromosomes. (C) Top 4 TF-MODISCO contribution weight matrix (CMW) motifs derived from count contribution scores of the human NSC ATAC ChromBPNet no bias model. (D) Selected TF-MODISCO CMW motifs identified in RGCs or IPCs from cortical organoids (in addition to those shown in Figure 5B).

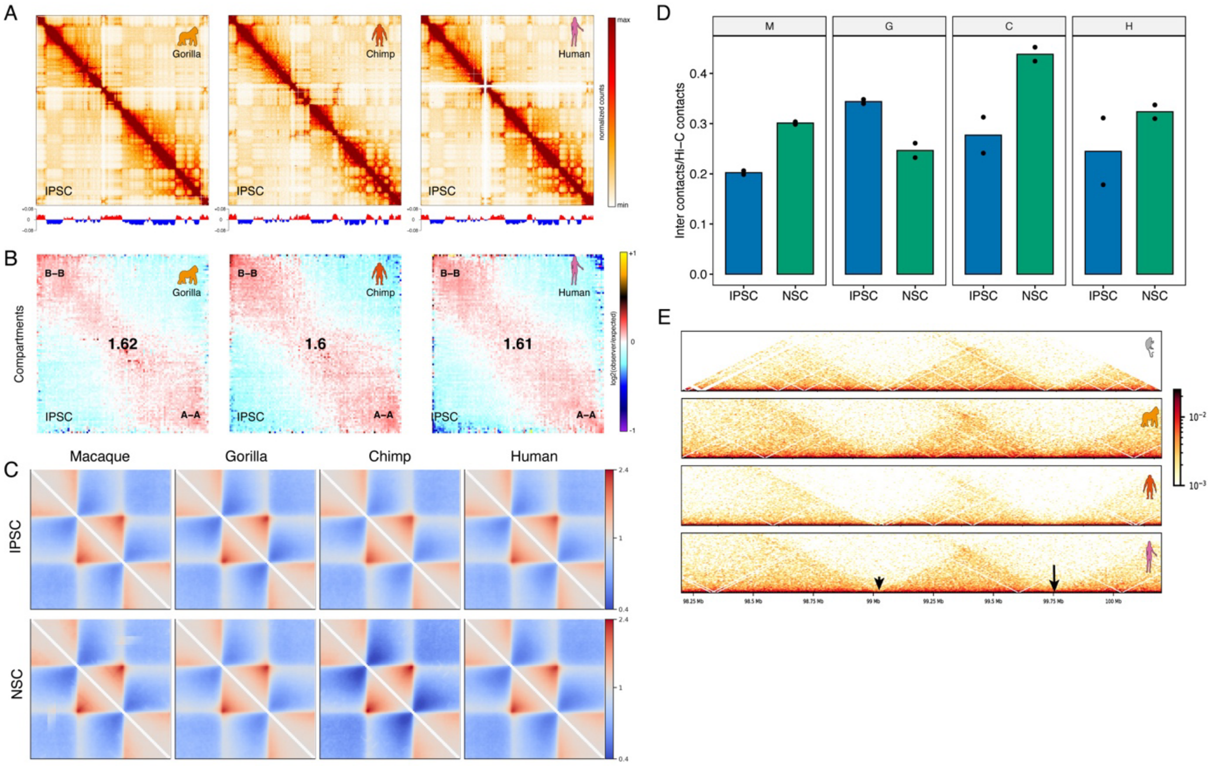


**Figure S6. Dynamics of the 3D genome during primate neocortex development and evolution**

(A) Observed contact matrices for chr3 at 250kb resolution (top) and the first eigenvector at 100kb resolution (bottom) for gorilla, chimpanzee, and human IPSC. (B) Saddle plots showing the average contact enrichment in IPSC at pairs of 100kb *loci* arranged by their eigenvalue. Numbers represent the compartment strength. (C) Average contact enrichment in IPSC (top) and NSC (bottom) across TADs. (D) Ratio of the inter-chromosomal to the total Hi-C contacts for primate IPSC and NSC. Dots represent individual measurements from each replicated experiment (n=2). (E) Normalized observed contact matrices at the *CNTN5 locus* for primate NSC. The previously identified conserved TAD boundary is marked by an arrowhead, while the arrow on the right highlights the reported human-specific TAD boundary[21]. Note that the insulation appears highly conserved across both regions.

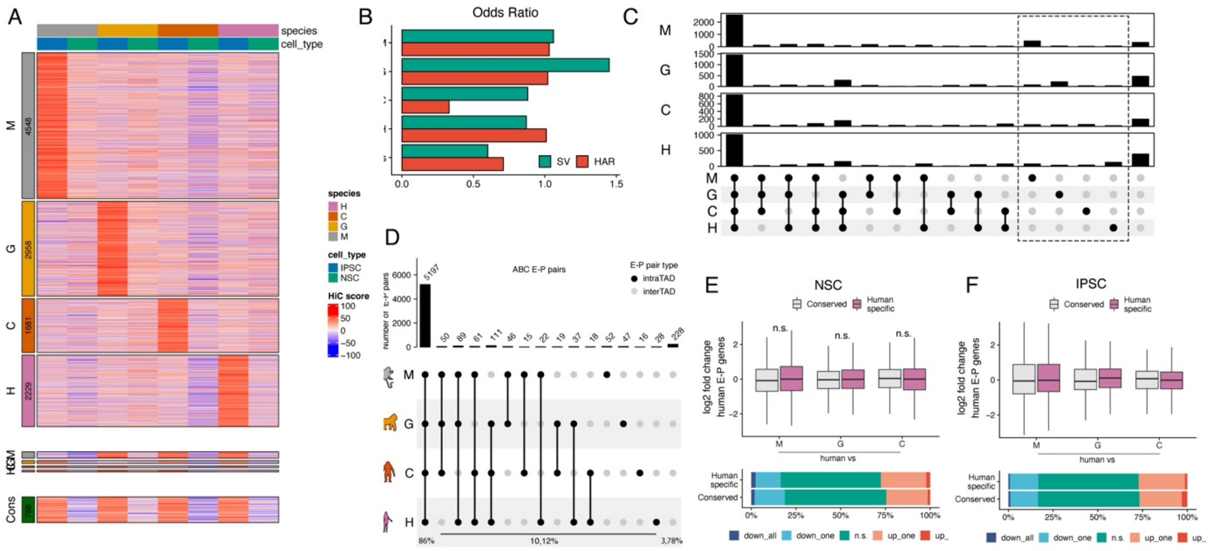


**Figure S7. Genome-wide comparative analysis of enhancer-promoter interactions**

(A) Heatmap depicting differential IPSC E-P interactions. (B) Odds ratio of the different categories of IPSC E-Ps overlapping with fhSVs[45] or HARs[23]. (C) Upset plot showing the intersection between different IPSC E-P classes and TADs in each species. The dashed rectangle indicates the interactions that are intra-TAD in one species but inter-TAD in all the other species. Note the increase of inter-TAD E-P interactions per group for each corresponding species. (D) Upset plot showing the intersection between E-P pairs based on the ABC model in human NSC and TADs in each species. (E) Boxplots displaying gene expression log2 fold change in human compared to NHP for IPSC human-specific and conserver E-P loops. Statistical significance was calculated using an unpaired two-sided Wilcoxon rank-sum test. (F-G) Stacked barplot showing the percentage of genes that are downregulated in all species (down_all), downregulated in only one species (down_one), not significant (n.s.), upregulated in one (up_one) or in all species (up_all) when comparing expression in human vs NHP NSC (F) or IPSC (G). M, crab-eating macaque; G, gorilla; C, chimpanzee; H, human; Cons, conserved.
